# Structural Basis of μ-Opioid Receptor-Targeting by a Nanobody Antagonist

**DOI:** 10.1101/2023.12.06.570395

**Authors:** Jun Yu, Amit Kumar, Xuefeng Zhang, Charlotte Martin, Pierre Raia, Antoine Koehl, Toon Laeremans, Jan Steyaert, Aashish Manglik, Steven Ballet, Andreas Boland, Miriam Stoeber

**Affiliations:** Department of Molecular and Cellular Biology, University of Geneva, Geneva, Switzerland; Department of Cell Physiology and Metabolism, University of Geneva, Geneva, Switzerland; Departments of Chemistry and Bioengineering Sciences, Vrije Universiteit Brussel, Brussels, Belgium; Department of Plant Sciences, University of Geneva, Geneva, Switzerland; Department of Electrical Engineering and Computer Sciences, University of California, Berkeley, CA, USA; Confo Therapeutics N.V., Gent, Belgium; Structural Biology Brussels, Vrije Universiteit Brussel, Brussels, Belgium; VIB-VUB Center for Structural Biology, VIB, Brussels, Belgium; Department of Pharmaceutical Chemistry, University of California, San Francisco, CA, USA

## Abstract

The μ-opioid receptor (μOR), a prototypical member of the G protein-coupled receptor (GPCR) family, is the molecular target of opioid analgesics such as morphine and fentanyl. Due to the limitations and severe side effects of currently available opioid drugs, there is considerable interest in developing novel modulators of μOR function. Most GPCR ligands today are small molecules, however biologics, including antibodies and nanobodies, are emerging as alternative therapeutics with clear advantages such as affinity and target selectivity. Here, we describe the nanobody NbE, which selectively binds to the μOR and acts as an antagonist. We functionally characterize NbE as an extracellular and genetically encoded µOR ligand and uncover the molecular basis for µOR antagonism by solving the cryo-EM structure of the NbE-µOR complex. NbE displays a unique ligand binding mode and achieves µOR selectivity by interactions with the orthosteric pocket and extracellular receptor loops. Based on a β-hairpin loop formed by NbE that deeply inserts into the µOR and centers most binding contacts, we design short peptide analogues that retain µOR antagonism. The work illustrates the potential of nanobodies to uniquely engage with GPCRs and describes novel μOR ligands that can serve as a basis for therapeutic developments.

## Introduction

G protein-coupled receptors (GPCRs) represent key therapeutic targets due to their central roles in cellular signaling and control over a plethora of physiological processes. Developing new ligands that bind a given GPCR with high selectivity remains a significant challenge in drug discovery (*1–3*). Small molecule ligands have historically dominated the landscape of GPCR-targeted drugs but recently biologics, including antibodies and nanobodies (Nbs), have emerged as an alternative class of ligands that offer distinct advantages and hold promise for therapeutic developments (*4*, *5*). Nbs are single domain antibody fragments derived from heavy chain-only antibodies, which naturally occur in camelids and cartilaginous fish, and are characterized by a small size, strong antigen-binding affinity, and binding loops that can access deep cavities on target proteins (*6*). Nbs can show enhanced selectivity over small molecules due to their ability to interact with unique and extended epitope surfaces. Over the last decade, Nbs that bind GPCRs on their intracellular side have served as innovative research tools to uncover GPCR signal transduction mechanisms (*7*, *8*). For example, Nbs were used as crystallization chaperones or as fiducial markers in high resolution structural studies (*9–12*). Conformation-selective Nbs were also repurposed into biosensors to report GPCR activity in living cells (*13*, *14*). Contrastingly, only a few Nbs that bind GPCRs as extracellular ligands and thereby modulate receptor function have been described (*15–18*). Generating knowledge on GPCR-targeting Nbs is key to unlocking their potential as both versatile research tools and therapeutic compounds.

Opioid receptors (ORs) are prototypical members of the rhodopsin-like GPCR family and function in pain modulation and analgesia (*19*, *20*). The OR family comprises four major receptor subtypes, including the μOR, δOR, κOR, and nociceptin-OR (NOPR), with the μOR representing the prime therapeutic target for pain relief. Approved drugs that target μORs are diverse small molecule compounds, including the widely used analgesics morphine and fentanyl. The ligand repertoire has recently been expanded through structure-based molecular docking, rational design, and high-throughput screening, delivering new OR ligands with distinct pharmacological profiles, including biased agonism, receptor subtype selectivity, and pharmacokinetic properties (*21–23*). Discovering additional and innovative modulators of μOR function, including agonists and antagonists, remains a pressing necessity for developing improved analgesics and compounds that can reduce or reverse the deleterious opioid side effects (*24*). Until now, no antibody or Nb ligand for ORs has been characterized in-depth, representing a hurdle in exploiting the unique features of biologics to effectively target ORs.

In this study, we functionally and structurally analyze NbE, an extracellular µOR-targeting Nb ligand (*9*). Using cellular binding and signaling assays, we identify that NbE selectively binds to μOR with nanomolar affinity and acts as an antagonist when added as ligand or expressed as a genetically-encoded cell surface displayed protein. We then solve the cryo-EM structure of the NbE-μOR complex, which reveals that NbE deeply inserts into the orthosteric pocket and additionally interacts with two extracellular loops (ECL) of the μOR. We find that the ECL interactions significantly contribute to NbE binding and OR subtype selectivity. Based on NbE’s complementary-determining region 3 (CDR3), which inserts as a β-hairpin loop into the orthosteric pocket, we synthesize peptide Nb mimetics of different lengths that retain binding and μOR-selective antagonism. The work uncovers a novel and unique ligand engagement profile at the μOR and provides a strategy for developing biologics-based µOR ligands.

## Results

### NbE binds the extracellular side of μOR and is an antagonist

We first tested whether NbE, which was part of a Nb library previously generated against the μOR (*9*), binds to the extracellular side of μOR in the surface of living cells. We covalently conjugated purified NbE with Alexa Fluor 488 and incubated OR-expressing HEK293 cells with the fluorescently labeled NbE. Confocal microscopy analyses showed a pronounced NbE signal at the plasma membrane of μOR-expressing cells after incubation with NbE (Fig. 1a). Cells expressing the closely related δOR, κOR, or NOPR were not labeled by NbE. Selective binding of NbE to μOR-expressing cells was also detected by flow cytometry analyses after adding NbE at increasing concentrations (Fig. 1b). δOR-, κOR-, or NOPR-expressing cells gated for similar receptor surface levels did not exhibit NbE binding above control cells (Fig. 1b). Since the NbE staining intensity of μOR expressing cells did not plateau at the highest NbE concentration tested (10 μM), we turned to grating-coupled interferometry (GCI) to determine the NbE binding strength. We immobilized biotinylated AVI-tagged NbE on a streptavidin coated biosensor surface and perfused it with buffer containing purified μOR at different concentrations. From the binding curves we extracted the affinity of the NbE-μOR interaction (K_D_=56 nM) and the kinetic parameters (k_a_=2.8*10^3^ M^-1^s^-1^ and k_d_=1.6*10^-4^ s^-1^), which revealed a slow NbE off-rate (Suppl. Fig. S1a).

**Figure 1:**
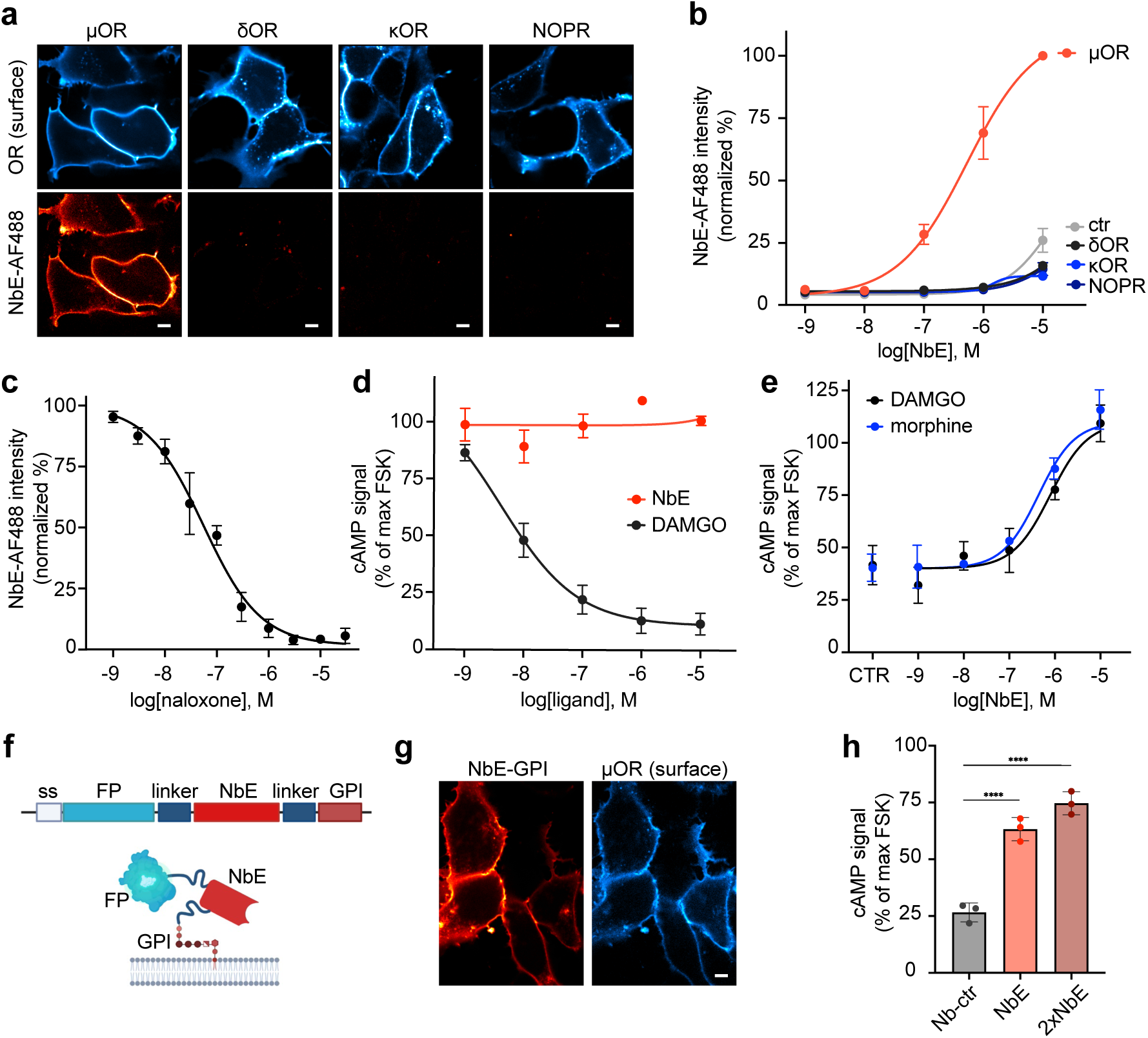
NbE binds the extracellular side of the μOR and acts as an antagonist. **(a)** Confocal images of HEK293 cells expressing FLAG-μOR, δOR, κOR, or NOPR (labeled with anti-FLAG M1-AF647, cyan) and incubated with 1 μM purified AF488-labeled NbE (red). Scale bar, 10 μm. **(b)** FACS-based quantification of AF488-NbE binding to HEK293 cells expressing μOR, δOR, κOR, or NOPR at different NbE concentrations (cells gated for similar receptor surface levels), N ≥ 3, mean ± SEM. **(c)** FACS-based quantification of AF488-NbE binding (1 μM) to μOR-expressing HEK293 cells pretreated with naloxone at different concentrations. N ≥ 3, mean ± SEM. **(d)** Maximum cAMP response in μOR- expressing HEK293, stimulated with 2.5 μM forskolin (FSK, norm. to 100%), treated with increasing concentrations of DAMGO or NbE. **(e)** Maximum cAMP response in μOR-expressing HEK293, stimulated with 2.5 μM FSK (norm. to 100%), treated with 10 nM DAMGO or 30 nM morphine (CTR) and pre-incubated with increasing concentrations of NbE. N ≥ 3, mean ± SEM. **(f)** Targeting NbE to the extracellular leaflet of the plasma membrane by fusion to a GPI anchor motif. ss = signal sequence, FP = fluorescent protein. **(g)** Confocal images of HEK293 cells stably expressing μOR (labeled with anti-FLAG M1-AF647, cyan) and transfected with mRuby2-NbE-GPI (red). Scale bar, 10 μm. **(h)** Maximum cAMP response in μOR-expressing HEK293, expressing GPI-anchored Nb-ctr, NbE, or bivalent NbE (2xNbE). All conditions treated with 10 nM DAMGO and normalized to cAMP level upon addition of 2.5 μM FSK. N = 3, mean ± SEM. ****P = <0.0001, by ordinary one-way ANOVA.

Next, we investigated if NbE binds competitively with orthosteric μOR ligands. We incubated μOR- expressing HEK293 cells with increasing concentrations of naloxone, subsequently added fluorescent NbE, and quantified the NbE signal by flow cytometry. NbE binding decreased in a naloxone concentration-dependent manner and was entirely abolished at high naloxone concentrations (Fig. 1c). The data suggested that the NbE binding site on the μOR overlaps with the orthosteric ligand binding pocket, and thus we tested if NbE modulates μOR activity. We first assessed whether NbE behaves as an agonist by measuring μOR-mediated inhibition of cyclic AMP (cAMP) accumulation in living cells, a readout of Gi-driven OR signaling. Application of NbE did not activate μOR even at high concentrations, in contrast to the peptide agonist DAMGO (Fig. 1d). We then tested whether NbE acts as a μOR antagonist and reduces the signaling effects of opioid peptides and small molecule opioid drugs. Indeed, pre-incubation of μOR-expressing cells with NbE caused a concentration-dependent decrease in DAMGO- and morphine-induced Gi signaling (Fig. 1e). At high concentrations, NbE fully blocked DAMGO and morphine-driven μOR inhibition of cAMP production. Furthermore, NbE addition strongly reduced the recruitment of a G protein probe (miniGi) and β-arrestin2 to DAMGO-activated μOR, as measured by split nanoluciferase (NanoLuc)-based complementation assays (Suppl. Fig. S1b).

We hypothesized that NbE may also antagonize μOR function when ectopically expressed in cells and targeted to the extracellular leaflet of the plasma membrane via a glycolipid anchor. To test this, we fused NbE to a C-terminal glycosylphosphatidylinositol (GPI) signal peptide, separated by a flexible linker (Fig. 1f). We also added a secretory signal peptide at the N-terminus, followed by an HA-tag or a fluorescent protein to enable determining the subcellular localization of the NbE fusion proteins (Fig. 1f). Confocal microscopy analyses of NbE-GPI-expressing cells showed that NbE predominantly localized at the cell surface (Fig. 1g). Immunostaining of cells expressing HA-NbE-GPI with HA antibodies in non-permeabilizing conditions revealed that NbE was efficiently displayed on the extracellular side of the plasma membrane (Suppl. Fig S1c). We also constructed a bivalent tandem NbE-NbE-GPI construct as well as a non-targeting Nb-GPI control (Nb-ctr), and both showed similar predominant cell surface localization (Suppl. Fig S1d). μOR-expressing HEK293 cells transfected with monovalent or bivalent NbE-GPI constructs showed significantly higher cAMP levels after DAMGO treatment relative to cells expressing the Nb-GPI control, indicating a NbE-mediated block of μOR signaling (Fig. 1h).

Taken together, NbE specifically binds to the extracellular side of the μOR and competes with orthosteric opioid ligands. Furthermore, NbE acts as an antagonist when added to cells as an extracellular ligand or when ectopically expressed as GPI-anchored fusion protein.

### Cryo-EM structure determination of the NbE-μOR complex

Advancement of structural biology techniques have led to the determination of several µOR structures in complex with either antagonists (*11*, *25*), partial agonists (*23*, *26*), or full agonists, including the endogenous peptides β-endorphin and endomorphin (*9*, *26–28*). All ligands bind to the orthosteric ligand binding pocket that is on the extracellular side and largely solvent exposed. To identify the precise molecular binding mode of NbE, we determined the structure of the NbE-µOR complex using cryo-EM. Initial structure determination was precluded due to the absence of large extracellular features of the complex. The surrounding detergent micelle additionally impaired accurate particle projection alignments, resulting in unsuccessful 3D reconstruction. To provide extra features for accurate particle alignment, we incubated the purified NbE-μOR complex with a Fab module consisting of a Nb-binding Fab fragment (NabFab) and an anti-Fab Nb, recently developed as a fiducial marker (*29*). The resulting stable and homogenous NbE-µOR-Fab module complex was purified by size exclusion chromatography (Suppl. Fig. S2a,b). Cryo-EM analyses enabled us to determine the structure of the NbE-µOR-Fab module complex at a global resolution of 3.1 Å (Suppl. Fig. S2c-f and Suppl. Fig. S3, Table 1). The EM-derived Coulomb potential map provided excellent side chain densities for most parts of the complex with the best resolved region at the NbE-µOR interface (Suppl. Fig. S2d,f).

The overall structure of the NbE-µOR complex bound to the Fab module is shown in Fig. 2a. The Fab fragment binds in a single and rigid conformation to NbE, enabling high-resolution structure determination of the NbE-µOR complex. For reasons of clarity, the Fab module is omitted in all subsequent figures (Fig. 2b and Suppl. Video 1). When bound to NbE, the µOR adopts an inactive conformation, which closely resembles the structures of the µOR bound to the morphinan antagonist β-funaltrexamine (β-FNA) or alvimopan (Suppl. Fig. S4a) (*11*, *25*). The average root mean square deviation (RMSD) for all C_α_ atoms of µOR between the NbE-µOR complex and the µOR complexes bound to antagonists is 0.78 Å, whereas the RMSD between NbE-µOR and agonist-bound structures is on average 1.72 Å (Suppl. Table 2). In particular, transmembrane helix 6 (TM6), which is displaced by roughly 10 Å in the activated state (*9*, *27*), superimposes well between the NbE-µOR and the β-FNA- or alvimopan-bound inactive µOR (Suppl. Fig. S4a,c). Moreover, a conserved core triad consisting of the amino acids I155^3.40^, P244^5.50^, and F289^6.44^

**Figure 2:**
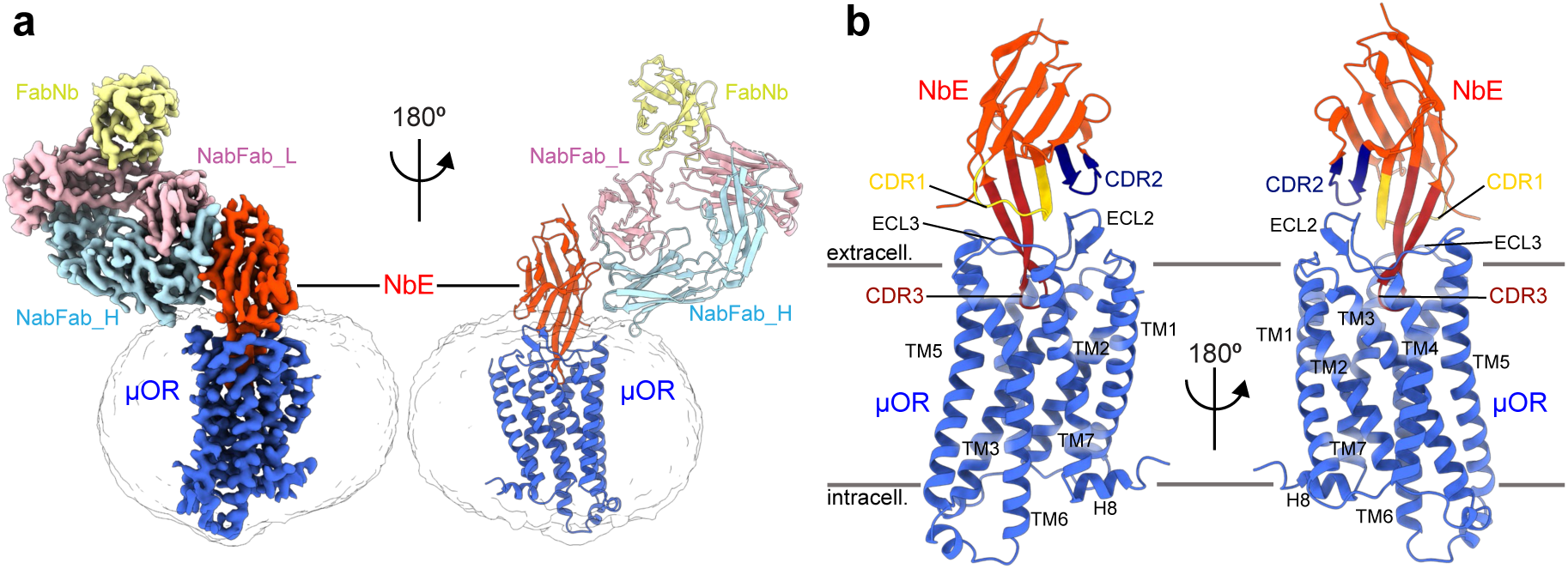
Architecture of the NbE-µOR complex. **(a)** Cryo-EM map (left) and ribbon representation (right) of the NbE-µOR-Fab module complex, with the µOR coloured in blue, NbE in red, the heavy and light chains of the Fab fragment in cyan and rose, respectively, and the anti-Fab nanobody in yellow **(b)** The NbE-µOR complex as ribbon representation, with a focus on the NbE-µOR interface. NbE deeply inserts its CDR3 β-hairpin loop into the orthosteric binding pocket. For reasons of clarity the NabFab module is omitted. CDR1-3 are highlighted in yellow, dark blue and dark red, respectively. Otherwise color-coded as in (a).

**Figure 3:**
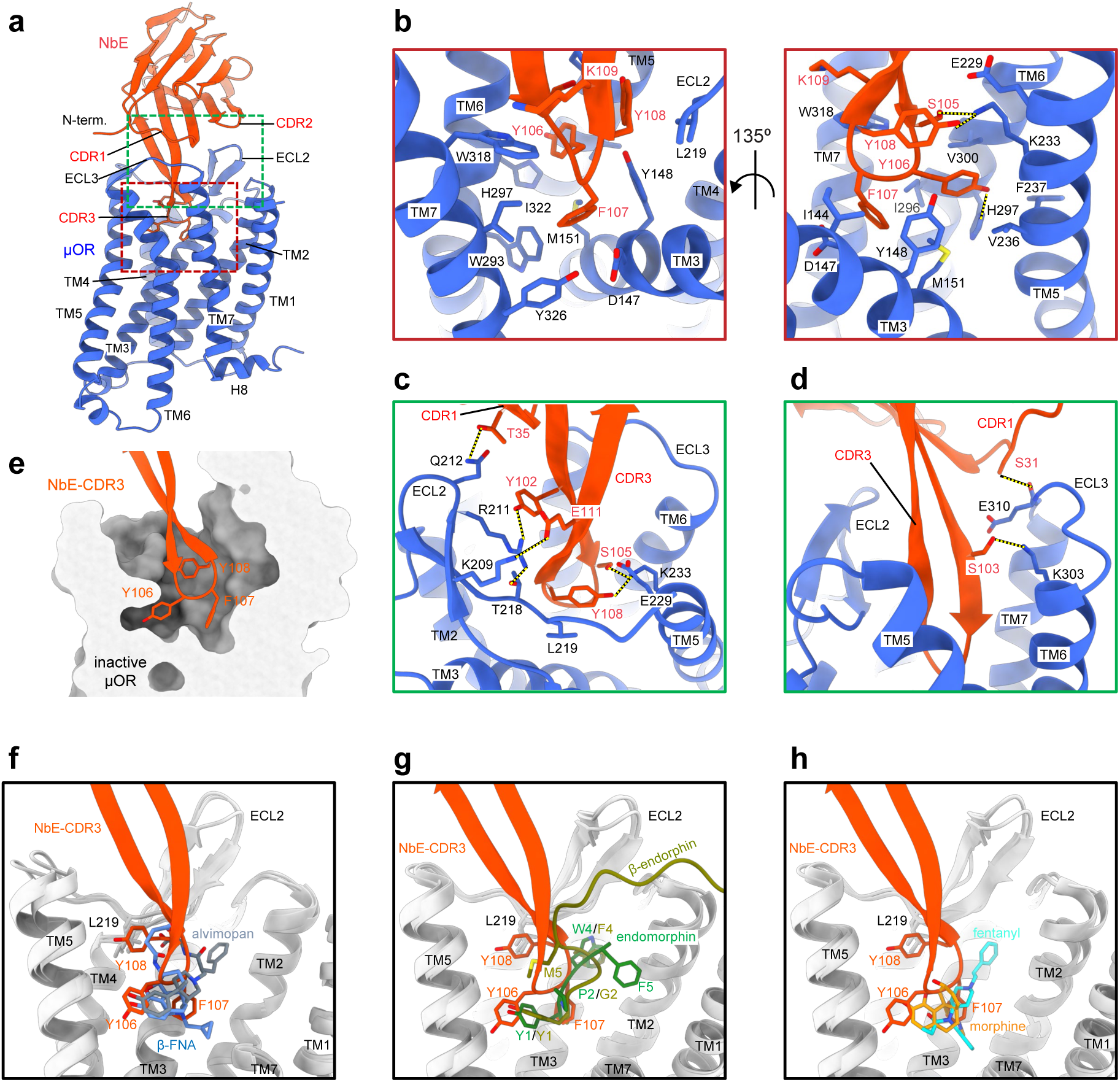
The NbE-µOR interaction interface. **(a)** Overview of the NbE-µOR complex, color-coded as in Fig. 2. Dashed squares in red and green indicate the close-up views shown in b-d. **(b)** Close-up of the orthosteric binding pocket, with interface residues shown as sticks. NbE residues F107 (left panel) and Y106 (right panel) are centered. Interface residues shown as sticks. **(c)** Top view of the µOR ligand binding pocket with CDR3^NbE^ deeply inserted. Interactions between ECL2 of the µOR and CDR3 of NbE are highlighted. Interface residues are shown as sticks and hydrogen bonds and salt bridges are indicated as yellow dashed lines. **(d)** Interaction interface between ECL3 of the µOR and CDR3 of NbE. Interface residues are shown as sticks and hydrogen bonds and salt bridges are indicated as yellow dashed lines. **(e)** The tip-forming residues Y106, F107, and Y108 (shown as sticks) of CDR3 are binding to distinct hydrophobic patches inside the orthosteric ligand binding pocket (shown as surface representation). **(f)** Binding mode superposition of NbE and the small molecule antagonists β-FNA (PDB: 4DKL) and alvimopan (PDB: 7UL4). NbE’s Y108 stacks against L219^ECL2^, which is a unique ligand-receptor interaction. Alvimopan inserts a phenyl group in a binding cleft not occupied by NbE or β-FNA. **(g)** Binding mode superposition of NbE and opioid peptide agonists β-endorphin (PDB: 8F7Q) or endomorphin (PDB: 8F7R). Y108^NbE^ uniquely recognises L219^ECL2^, whereas a phenol group of endomorphin binds to a similar site also recognised by alvimopan (f) or fentanyl (h). **(h)** Binding mode superposition of NbE and the small molecule agonists fentanyl (PDB: 8EF5) or morphine (PDB: 8EF6).

(Ballesteros-Weinstein numbering scheme (*30*)), which lies below the ligand binding pocket and propagates structural rearrangements involved in receptor activation (*9*), superimposes well with the µOR in the inactive form (Suppl. Fig. S4b,d). Structural differences in the ligand binding pocket between agonists, partial agonists, and antagonists are relatively subtle (*31*, *32*), however, binding of NbE to the µOR induces several unique conformational changes in the orthosteric binding pocket and the extracellular loops.

### Unique interaction profile of NbE with the inactive μOR

Binding of NbE to the µOR is mediated through its complementarity-determining regions (CDRs) CDR1 and CDR3, with the main interaction interface being formed between the β-hairpin loop of CDR3 and µOR’s TM helices 3, 5, 6, and 7 (Fig 3a-d, Suppl. Table 3). CDR3 deeply inserts into the orthosteric ligand binding pocket with the three aromatic residues Y106^NbE^, F107^NbE^ and Y108^NbE^ forming the tip of the loop (Fig. 3b,e, Suppl. Video 1).

Similar to previously determined µOR structures bound to the ligands DAMGO (*27*), BU72 (*9*), β-FNA (*25*), alvimopan (*11*), PZM21, and FH210 (*23*), as well as β-endorphin and endomorphin (*28*), H297^6.52^ is positioned closely to a phenol hydroxyl group of the NbE ligand (Fig. 3b). While binding of BU72 and β-FNA is mediated through a hydrogen bond network that involves two water molecules (*9*, *25*), in the NbE-µOR complex, H297^6.52^ and Y106^NbE^ are directly forming a hydrogen bond (Fig. 3b). In addition, Y106^NbE^ is surrounded by mainly hydrophobic residues, including Y148^3.33^, M151^3.36^, V236^5.42^, F237^5.43^, I296^6.51^, and V300^6.55^ (Fig. 3b). F107^NbE^, the second tip-forming aromatic residue, inserts itself into a neighboring hydrophobic cavity (Fig. 3b,e) and interacts with the key residues I143^3.30^, Y148^3.33^, M151^3.36^, W293^6.48^, I322^7.39^ and Y326^7.43^ of µOR (Fig. 3b). Interestingly, the aspartate D147^3.32^, present in many family A GPCRs, and crucial for the recognition of DAMGO, β-FNA, BU72, PZM21, and endogenous peptides by forming a salt bridge with an amine group of the ligand, is rotated compared to all other µOR structures (Suppl. Fig. S4e). Instead of forming a salt bridge with the ligand, in this conformation D147^3.32^ stacks onto F107^NbE^, representing a novel ligand-receptor interaction mode (Fig. 3b and Suppl. Fig. S4e). The third aromatic residue Y108^NbE^ is situated at the periphery of the orthosteric binding pocket (Fig. 3c). The aromatic ring of the phenol group stacks against L219^ECL2^, part of ECL2, whereas the hydroxyl group forms a salt bridge with K233^5.39^ (Fig. 3b,c). K233^5.39^ had previously been identified as the side chain to which the morphinan ligand β-FNA is covalently attached (*25*, *33*). Another notable difference in the ligand binding pocket involves W318^7.35^ that is uniquely positioned likely due to steric constraints when inserting the large NbE ligand into the narrow ligand binding pocket (Suppl. Fig. S4e). This specific rotamer conformation has only been observed when DAMGO, a pentameric peptide ligand, is bound to the µOR, however, in the NbE-µOR structure W318^7.35^ is shifted by several Å. Because of the unusual positioning of W318^7.35^ and its rotamer conformation, K303^6.58^ that often stacks onto W318^7.35^ is displaced by several Å, forming a salt bridge with S103^NbE^ of NbE instead (Fig. 3d).

The three tip-forming aromatic residues of NbE are recognized by neighboring but distinct binding cavities in the orthosteric binding pocket (Fig. 3e). Here, Y106^NbE^ and F107^NbE^ are binding to sites usually occupied by small molecule ligands (Fig. 3f-h), whereas Y108^NbE^ stacks onto L219 of ECL2, thereby stabilizing the CDR3 β-hairpin loop in its conformation. Binding of ligands to this peripheral binding site has not yet been observed in other µOR structures. Conversely, the ligands alvimopan (antagonist), endomorphin, and fentanyl (agonists) are inserting aromatic functional groups into another binding cavity, which is not occupied by NbE (Fig. 3f-h).

To probe if the Nb ligand undergoes conformational rearrangements upon µOR binding, we crystallized NbE in its unbound form. The crystal contained three NbE molecules per asymmetric unit. All three molecules are virtually identical to each other, including the CDRs. Comparing the unbound and bound NbE structures revealed that CDR1 and the CDR3 β-hairpin loop undergo a conformational shift when µOR-bound (Suppl. Fig. S5a,b). The conformational differences imply that CDR1 and CDR3 exhibit intrinsic flexibility, potentially important for conformational selection of the inactive µOR and high-affinity binding.

In summary, NbE shows a unique interaction profile with the µOR. In particular, D147^3.32^, L219^ECL2^, and W318^7.35^ of the µOR engage with NbE in a so far undetected ligand binding mode that can offer new possibilities for structure-guided drug design.

### Extracellular loops confer NbE binding selectivity

To provide a rationale for NbE’s selectivity for µOR over other OR family members (Fig. 1a,b), we focused our structural analyses on the extracellular loops, which represent the most variable regions in the OR family. In particular, ECL2 shows variations in sequence and length between the receptors (Fig. 4a, Suppl. Fig. S6), providing a possible explanation for the observed µOR selectivity given the specific interactions of NbE with ECL2 and ECL3 (Fig. 3c,d). K209^ECL2^, R211^ECL2^ and Q212^ECL2^ form salt bridges with E111^NbE^, Y102^NbE^, and T35^NbE^ respectively, thereby contributing to high-affinity binding (Fig. 4b). Importantly, Q212^ECL2^ is unique to the µOR. The centered L219^ECL2^ of ECL2 stacks onto Y108^NbE^ of CDR3 (Fig. 3b). K303^6.58^, also unique to µOR, forms a specific salt bridge with S103^NbE^ that likely stabilizes ECL3 in a conformation that allows several nonspecific side chain-main chain interactions between µOR and NbE, with E310^ECL3^ being at ECL3’s center (Fig. 4c). Binding of NbE to µOR induces clear shifts for ECL2, parts of TM6, ECL3 and TM7 compared to the other two antagonist-bound µOR structures (Suppl. Fig. S4a).

**Figure 4:**
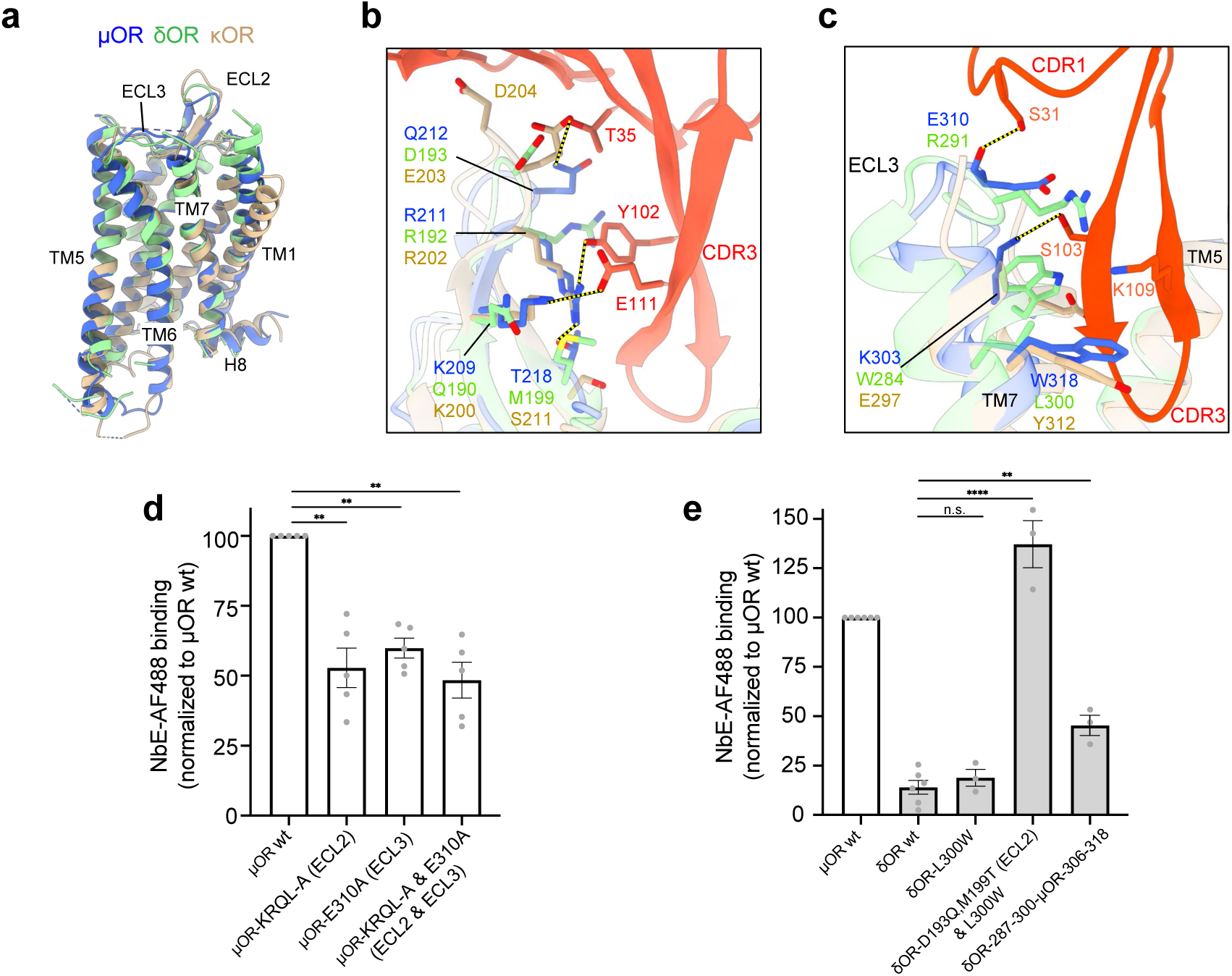
ECL regions confer NbE binding selectivity. **(a)** Superposition of the inactive state of the NbE-bound µOR (blue), inactive δOR (green, PDB: 4EJ4), and inactive κOR (brown, PDB: 4DJH) with ECL2 and ECL3 indicated. **(b)** Binding interface of different ECL2 regions (µOR, δOR, and κOR) with NbE. For comparison, the δOR and κOR have been superimposed onto the µOR. Q212^ECL2^ is unique to the µOR. **(c)** Binding interface of TM7 and ECL3 regions from the µOR, δOR, and κOR with NbE. For comparison, the δOR and κOR have been superimposed onto the TM7 and ECL3 regions of the µOR. Most interface residues differ between the different receptor subtypes. Color-coded as in (a). **(d)** NbE-binding to the wild-type (wt) µOR and its ECL variants. NbE binding to wt µOR was normalized to 100%. µOR-KRQL-A mutant: K209^ECL2^, R211^ECL2^, Q212^ECL2^ and L219^ECL2^ substituted by alanine, E310A mutant: E310^ECL3^ substituted by alanine. µOR-KRQL-A & E310A: both sets of mutations combined. N = 3, mean ± SEM. **P = 0.0012 (ECL2), 0.0047 (ECL3), 0.0010 (ECL2&3) by ordinary one-way ANOVA. **(e)** NbE binding to δOR mutants. As in (d), NbE-binding to wt µOR was normalized to 100%. The three δOR mutants include δOR L300W^7.35^, a triple δOR^D193Q, M199T, L300W^ mutant, and a δOR mutant with the entire ECL3 and distal parts of TM7 substituted by µOR residues (δOR^287-300-µOR-306-318^). N ≥ 3, mean ± SEM. **P = 0.0011, ****P = <0.0001, n.s. = 0.8998 by ordinary one-way ANOVA.

To probe the relevance of µOR’s ECLs in NbE binding, we first substituted ECL2 residues K209^ECL2^, R211^ECL2^, Q212^ECL2^ and L219^ECL2^ (‘KRQL’ motif) by alanine and tested NbE binding to the µOR mutant by flow cytometry. Compared to wild-type µOR, the KRQL-A mutant showed 50% reduced NbE binding, indicating a lower affinity for the ECL2 mutant (Fig. 4d). Exchanging E310^ECL3^ in CDR3 with alanine also significantly reduced NbE binding (Fig. 4d). Combining both the ECL2 and ECL3 mutations did not lead to further reduction in NbE binding, highlighting the central role of the orthosteric pocket interactions (Fig. 4d).

Next, we reasoned that assimilating the δOR to the µOR, based on the uncovered interaction interfaces, might transform the δOR from a non-binder to a NbE-binder. Given the established role of the amino acid position 7.35 in opioid receptor subtype selectivity (*34*, *35*), we first mutated δOR’s L300^7.35^ in the orthosteric pocket to the corresponding µOR residue W318^7.35^ (δOR^L300W^ mutant). We then also converted the δOR residues D193^ECL2^ (corresponds to µOR Q212^ECL2^) and M199^ECL2^ (corresponds to µOR T218^ECL2^) into glutamine and threonine residues respectively (Fig. 4b). T218^ECL2^ of µOR forms an intramolecular salt bridge with the guanidinium group of R211^ECL2^. As a consequence of this loop stabilization, K209^ECL2^ and R211^ECL2^ of the µOR are ideally positioned to create the aforementioned hydrogen bonding network with Y102^NbE^ and E111^NbE^ of NbE (Fig. 4b). Because T218^ECL2^ is unique to the µOR, we speculated that mutating δOR M199^ECL2^ into threonine might stabilize the ECL2 of the δOR in a confirmation that favors NbE binding. We quantified fluorescent NbE binding to cells expressing µOR (control), wild-type δOR, or δOR mutants by flow cytometry and gated cells for similar receptor levels. As expected, only non-specific background NbE binding was detected for cells expressing wild-type δOR (Fig. 4e). The NbE signal was not increased for cells expressing δOR^L300W^. However, strong NbE binding was observed when cells expressed the triple mutant δOR^D193Q, M199T, L300W^ (Fig. 4e). The results show that mutating two residues in the ECL2^δOR^, combined with the orthosteric L300W mutation, can convert the non-binding δOR into a strong NbE binder. To test the role of ECL3, we next substituted the entire ECL3 as well as residues of the connecting α-helix 7 of the δOR (residues 287-300) with residues of the µOR (306–318). The substitution led to a moderate but significant increase in NbE binding (Fig. 4e), indicating that µOR’s ECL3 region partially contributes to NbE binding.

In sum, the mutational and gain-of-function studies identify ECL2 and ECL3 as important contributors to NbE binding and receptor subtype selectivity.

### Peptide mimetics of NbE’s CDR3 bind and antagonize µOR

The centering of key contacts on a single CDR made NbE’s CDR3 a promising starting point in the design of ligands that downsize the Nb towards smaller peptides based on the antigen-binding paratope. We designed and synthesized a library of linear peptides of increasing length, with the shortest peptide based on the four residues ^105^SYFY^108^ that compose the β-turn segment at the tip of NbE’s CDR3 (Fig. 5a). We systematically extended each peptide by one N- and one C-terminal residue of the CDR3^NbE^ with the longest peptide spanning the 14 residues ^100^KYYSGSYFYKSEYD^113^ (Fig. 5a). Peptide binding to µOR was assessed using the HTRF Tag-lite binding assay, which relies on FRET between SNAP-tagged µOR labeled with terbium cryptate as FRET donor, and the red fluorescent opioid ligand naltrexone, serving as acceptor, with a decrease in FRET indicating competitive binding of a test compound. Binding of the control ligand naloxone and of NbE was readily detected with IC_50_ values of 14 nM and 40 nM respectively (Fig. 5b). In the Tag-lite assay, we detected significant binding of peptides 5 (12 residues) and 6 (14 residues) at IC_50_ values of 4.3 µM and 29 µM (Fig. 5b). The shorter peptides showed little to no binding (Suppl. Fig. S7). We next tested the ability of each peptide to antagonize DAMGO-driven µOR signaling by probing the cAMP response. Peptides 1-4 did not reverse the DAMGO effect, while peptides 5 and 6 caused a strong and concentration-dependent reduction of µOR signaling, detected as significantly increased intracellular cAMP levels (Fig. 5c). Furthermore, DPDPE-driven δOR signaling was not antagonized by peptide 5 at the tested concentrations, showing that the CDR3 peptide mimetic retains receptor subtype specificity (Fig. 5d). Taken together, the observed extensive interaction interface between the CDR3 of NbE and the µOR allowed the design of downsized linear peptide mimetics that retain µOR-specific antagonism.

**Figure 5:**
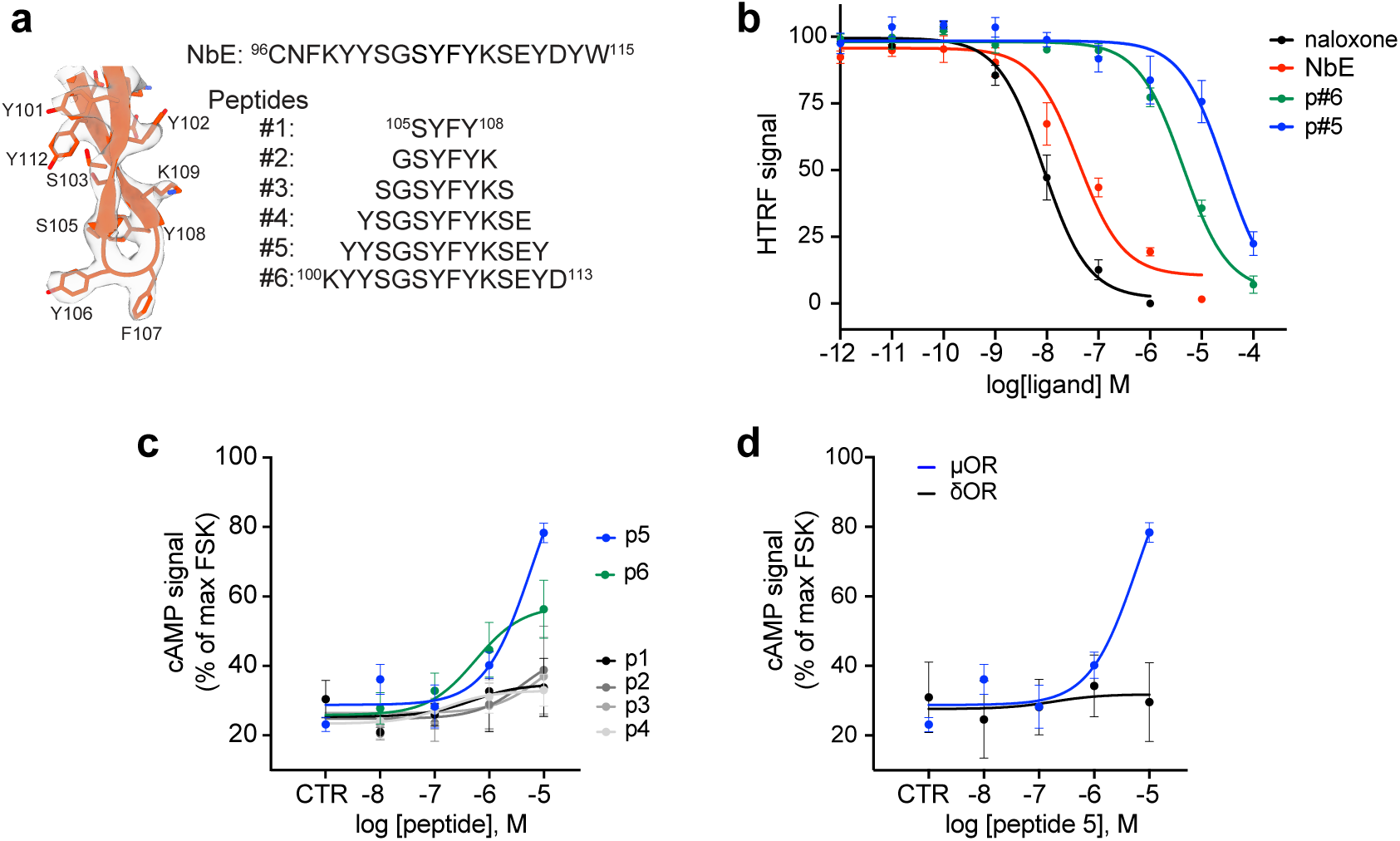
CDR3-based peptide-mimetics exhibit µOR binding and antagonism. **(a)** CDR3 conformation of NbE when bound to µOR (left), and peptides synthesized based on CDR3^NbE^ (#1 - #6). Peptides #1 - #5 are *N*-acetylated, *C*-terminal carboxamides, peptide #6 is an unacetylated *C*-terminal carboxylic acid for improved solubility. **(b)** HTRF competition binding assay profiles of naloxone, NbE, and peptides #5 and #6 of HEK293 cells stably expressing SNAP-μOR, labeled with SNAP-Lumi4-Tb and 3nM red-labeled naltrexone derivative, incubated with increasing concentrations of ligands. Normalized to cells without added compound; N ≥ 3, mean ± SEM. **(c)** Maximum cAMP response in μOR-expressing HEK293, stimulated with 2.5 μM FSK (norm. to 100%), treated with 10 nM DAMGO (CTR) and pre-incubated with increasing concentrations of peptides #1 - #6. N ≥ 3, mean ± SEM. N ≥ 3, mean ± SEM. **(d)** Maximum cAMP response of HEK293 cells stably expressing μOR or δOR, stimulated with 2.5 μM FSK (norm. to 100%), treated with 10 nM DAMGO or 1 nM DPDPE respectively (CTR), and pre-incubated with increasing concentrations of peptide #5. N ≥ 3, mean ± SEM.

The NbE-based paratope mimetics provide a new source of small peptide opioid ligands and demonstrate the value of Nbs for biologics-based drug development.

## Discussion

The emergence of biologics as therapeutic agents has opened new avenues in GPCR drug discovery (*4*). In this study, we present an in-depth molecular characterization of the single-domain antibody fragment NbE that functionally interacts with the µOR. NbE selectively binds to the extracellular domains of µOR with nanomolar affinity and antagonizes its function. The cryo-EM structure of the NbE-μOR complex shows that NbE stabilizes the µOR in an inactive conformation that resembles µOR structures bound to the small molecule antagonists β-FNA or alvimopan (*11*, *25*). Yet NbE exerts distinct effects on key residues in the orthosteric binding pocket as well as the extracellular regions of µOR. The interactions of NbE with µOR’s ECL2 and ECL3 critically contribute to selective binding, paralleling recent observations for the role of extracellular vestibules in serving as selectivity filters of ORs for different opioid peptide agonists (*28*). NbE exhibits a new ligand interaction profile with the µOR, centering almost all contacts on a hairpin loop formed by CDR3. The work opens the possibility to therapeutically target µOR with a biologic and provides an important structural template for new ligand design.

NbE’s binding kinetics are characterized by a relatively slow on-rate and by nearly irreversible binding on the time scale studied, pointing to unique bimolecular recognition. Based on the conformational differences that we detect in NbE’s unbound and µOR-bound state, in particular with regards to CDR3, it is possible that µOR binding drives NbE towards a conformation that is more complementary to the orthosteric pocket. Moreover, since GPCRs are known to be inherently dynamic and sample an ensemble of conformations in the unliganded state, NbE’s slow on-rate may also reflect conformational selection of a weakly populated µOR conformation. The slow off-rate makes NbE a unique µOR ligand, holding the potential for long-lasting µOR antagonism and distinct physiological effects.

Recent structures of class A GPCRs bound to extracellular Nbs begin to reveal the diversity in Nb driven GPCR modulation (Suppl. Fig. S8). For example, Nbs targeting the α1A-adrenergic receptor (*36*), rhodopsin (*18*), or orexin receptor 2 (*37*) interact with extracellular loops and allosterically modulate receptor function without directly contacting the orthosteric pocket. In contrast, extracellular Nbs of the apelin receptor (APJ) (*17*) or the angiotensin II type I receptor (AT1R) (*38*) occupy the orthosteric pocket and mimic the binding of a peptide ligand. Similar to the NbE-µOR interaction, the APJ and AT1R Nbs access the orthosteric pocket via their long CDR3 loops. Yet APJ and AT1R Nbs both exhibit extensive additional interactions via CDR1 and CDR2, which contrasts with the limited interactions outside the CDR3 region detected for NbE. We find that linear CDR3 peptide analogues retain biological activity and are selective for the µOR, confirming the importance of the CDR3 sequence in NbE binding and, more broadly, demonstrating that CDR paratopes of Nbs can spur the development of low molecular weight antibody mimetics. Until now, OR peptide antagonists have been developed through structural modification and conversion of natural agonist peptides or by functional screening of synthetic short peptide libraries, often yielding in peptides with limited OR subtype selectivity or reduced affinity (*39–41*). The functional NbE-derived peptides described here comprise the NbE CDR3 residues that interact with the orthosteric pocket and include flanking Y102 and E111 that form salt bridges with µOR’s ECL2, providing a rationale for their µOR selectivity. As expected, taking NbE’s CDR3 sequence out of the rigid Nb framework results in a drop in affinity and efficacy, however, future steps towards fine-tuning the three-dimensional peptide structure, for example via side chain-to-side chain macrocyclization or insertion of conformationally constrained amino acids, and will very likely give access to improved binding and antagonist potency.

Beyond NbE’s action as extracellular ligand, we present a strategy to employ NbE as a genetically encoded tool to modulate OR activity. Linkage of NbE to a GPI-anchor effectively displays NbE on the extracellular plasma membrane leaflet and blocks OR function, implementing novel ways for inhibiting OR function in neurons or sub-neuronal compartments in the future.

In sum, the present results reinforce the emerging notion that Nbs can uniquely engage with clinically important target proteins and constitute a new class of GPCR ligands. Mechanistic insights into Nb-GPCR engagement can instruct novel targeting strategies poised to impact future drug development.

## Material and Methods

### Mammalian cell culture conditions

HEK293 cells (CRL-1573, ATCC, female) were cultured in Dulbecco’s modified Eagle’s medium (DMEM, Gibco), supplemented with 10% fetal bovine serum (FBS, Thermo Fisher). HEK293 cells stably expressing N-terminally signal sequence FLAG (ssf)-tagged µOR (HEK293-µOR) were cultured in the presence of 250 μg/ml Geneticin (Gibco). For transient DNA expression, Lipofectamine 2000 (Invitrogen) was used according to the manufacturer’s instructions.

### NbE purification and labeling

*Escherichia coli* WK6 cells were transformed with NbE (originally named ‘Nb35’ (*9*)) cloned into the pXAP100 plasmid, containing an N-terminal pelB signal sequence for periplasmic expression and C-terminal 6xHis-tag for purification. A transformed colony was then grown in Terrific Broth media (1.2% w/v Tryptone, 2.4% w/v Yeast Extract, 0.6% Glycerol, 0.1% Glucose, 2mM MgCl2) at 37°C with 50 μg/mL ampicillin to an optical density (OD) at 600 nm of 0.7, followed by induction with 1 mM Isopropyl β-D-1-thiogalactopyranoside (IPTG, Biosolve Chimie, Catalog number 0010006204) and incubated overnight at 37°C for expression at 180 rpm shaking. Cell pellets were resuspended in TES buffer (0.2M Tris, 5mM EDTA, 0.5M sucrose, pH 8.0) equivalent to 5% of bacterial culture volume, and kept on a shaker at 4°C for at least 60 min. Cells then underwent an osmotic shock by adding double the volume of TES diluted 1:4 with H_2_O and kept on the 4°C shaker for 45 minutes. Cells were then pelleted and discarded, and NbE contained in periplasmic lysate purified by Ni^2+^-affinity (HisPur Ni NTA Resin; Thermo Fisher Scientific, Catalog number 88221) and eluted by low pH by adding Acetate Buffer (50 mM NaAc,1M NaCl, pH4.5-4.7) and immediately adding 1M Tris (pH 7.5) to the eluates. NbE was further purified by size-exclusion chromatography (Superdex 75, Cytiva) and purity assessed by SDS-PAGE gel electrophoresis and gel staining with Coomassie blue (Bio-Rad, Catalog number 1610400). NbE was covalently conjugated at primary amines of lysines with Alexa Fluor™ 488 (Invitrogen, Catalog number A10235). For conjugation, 1 mg NbE was incubated with AF488 for 1 hour at room temperature (RT) and AF488-NbE separated from free dye using gel filtration. The concentration and degree of labeling was determined with a NanoDrop spectrophotometer (Thermo Scientific) and AF488-NbE subsequently used in fluorescence microscopy- and flow cytometry-based binding assays.

### Flow cytometry-based binding assay

HEK293 cells were transiently transfected with ssf-tagged murine µOR, δOR, κOR, and NOPR or with µOR and δOR mutants. 24 h post transfection, cells were detached with PBS-EDTA and resuspended in ice-cold assay buffer (PBS, 1 mM Ca^2+^, 0.5 mM Mg^2+^) at 8 million live cells/ml. Cells were incubated for 30 min at RT with AF488-NbE at various concentrations and 10 minutes with AF647-conjugated anti-FLAG M1 antibody. Cells were washed twice with PBS and resuspended in FACS buffer (PBS, 1 mM Ca^2+^, 0.5 mM Mg^2+^, BSA 0.5%). Flow cytometry was performed using a Beckman Coulter CytoFLEX Flow Cytometer. Data were analyzed with FlowJoTM v10 software and cells were gated for singlets, living cells, and OR expression using the AF647 signal. To probe competitive binding with naloxone, HEK293-µOR cells were seeded at 5 × 10^^^4 live cells/cm^2^ density. Cells were detached and resuspended in ice-cold assay buffer. Cells were then pre-treated with different concentrations of naloxone for 5 minutes, followed by addition of 1 μM AF488-NbE for 30 minutes. Cells were washed twice with PBS, resuspended in FACS buffer, and analyzed using the Beckman Coulter CytoFLEX Flow Cytometer. Data were analyzed with FlowJoTM v10 software and cells were gated for singlets and further gated for similar receptor expression levels. Individual conditions were normalized to the maximum AF488 signal from the cells treated with 10 μM NbE-AF488.

### Luminescence-based cAMP assay

HEK293-µOR cells were transfected with a plasmid encoding pGloSensor-20F cAMP reporter (Promega). Cells were harvested 24 h post transfection and resuspended at a 1.5x10^^^6 live cells/ml in assay media (DMEM without phenol red or FluoroBrite DMEM, 250 μg/ml luciferin). 100 μl cell suspension was plated into each well of a clear bottom 96-well plate, and equilibrated for 60 min at 37°C. To probe the reversal of DAMGO or morphine effects, cells were pre-incubated with different concentrations of NbE. Before addition of ligands, sequential luminescence images were collected to obtain basal luminescence values using the FDSS/μCELL kinetic plate imager (Hamamatsu) with an integrated simultaneous dispensing head and simultaneous detection across the plate. Cells were then treated with ligands (10 nM DAMGO, 30 nM morphine) and 2.5 μM Forskolin (FSK, Sigma-Aldrich F6886), followed by continuous luminescence imaging for 10 min, using two technical replicates per condition. Luminiscence signal from the cells without Luciferin was considered as background. Difference in luminescence signals between the baseline and maximum signal was used for analysis. Cells treated with 2.5 μM FSK were used as maximum for normalization.

### Confocal microscopy-based binding assay

HEK293 cells were seeded on poly-L-lysine-coated 35-mm Cellvis glass-bottomed dishes (IBL, 220.110.022) and, after 24 hours, transfected with ssf-µOR, -δOR, -κOR, or -NOPR (0.8 μg DNA) using 3 μl of Lipofectamine 2000. 16 to 24 h post-transfection, cells were incubated at 37°C with 1 μM AF488-NbE for 30 minutes and Alexa Fluor™ 647 conjugated (Invitrogen, Catalog number A20173) M1 antibody for 10 min in HBS imaging solution (Hepes-buffered saline with 135 mM NaCl, 5 mM KCl, 0.4 mM MgCl2,1.8 mM CaCl2, 20 mM Hepes, 1 mM d-glucose, 1% FBS, adjusted to pH 7.4). Cells were subsequently washed 2x with PBS and imaged with a spinning disk confocal microscope (Nipkow, Zeiss) using Plan Apo 63x/1.4 Oil DICIII objective in a temperature and CO_2_-controlled environment (37°C, 5% CO2).

### Tag-lite HTRF binding assay

HEK293 cells stably expressing SNAP-µOR were labeled with 100 nM Tag-lite SNAP-Lumi4-Tb (Revvity Catalog number SSNPTBC). The labeled cells were resuspended at a concentration of 1x10^6 cells/ml. 10 µl of the resuspension was plated in each well of low volume 96-well plates or 384-well plates to obtain 10’000 cells per well. 5 µl of labeled antagonist (Revvity Catalog number L0005RED) was added to each well, resulting in a final concentration of 3 nM, followed by addition of different concentrations of the test compound, and incubation for 3 h at room temperature, protected from light. Each experimental condition was performed in technical triplicates. Upon reaching equilibrium after 3 h, the FRET signal at 620 nm and 660 nm was read using a SpectraMax Paradigm Multi-Mode Microplate Reader (Molecular Devices). The ratio of the acceptor and donor emission signals (HTRF ratio signal) for each individual well was calculated. HTRF signals were plotted against concentrations. All conditions were normalized to the maximum HTRF ratio from the wells with no labeled ligand. The signal from 100 nM naloxone condition was considered as background.

### Grating coupled interferometry assay

Grating Coupled Interferometry (GCI) experiments were conducted on a Creoptix WAVE delta system using 4PCP chips (Creoptix AG). The chips were conditioned with 100 mM sodium borate (pH 9.5) and 1 M NaCl (Xantec). Neutravidin (100 μg/μl in 10 mM sodium acetate, pH 5.0) was immobilized on the chip surface using standard amine coupling. This included 420 s of surface activation with a 1:1 mix of 400 mM EDC and 100 mM NHS (Xantec), 420 s neutravidin injection, 420 s BSA (0.5%) injection, and a final 420 s surface passivation with 1 M ethanolamine at pH 8.0. Subsequently, biotin-NbE was captured at three concentrations: 100 μg/μl, 20 μg/μl, and 1 μg/μl, yielding surface masses of approximately 3700, 3500, and 2300 pg/mm², respectively. Any remaining neutravidin sites were filled with 100 μg/μl biotin-BSA. All preparation steps were performed at 10 μl/min flow rate.

Analyte µOR was injected in a 1:2 dilution series from 7.8 nM to 1000 nM at 45 μl/min. The running buffer contained 20 mM HEPES (pH 7.5), 100 mM NaCl, 0.01% LMNG, and 0.002% CHS. Blank injections and a reference channel were used for double referencing, and a 0-2% DMSO calibration curve was employed for bulk refractive index correction.

Data were analyzed with Creoptix WAVEcontrol software, with corrections made for X and Y offset, DMSO calibration, and double referencing. A 1:1 Langmuir binding model with bulk correction was used for all experiments. Consistent results were observed across the three NbE concentrations.

### Purification of the anti-Fab Nb

A fragment encoding the anti-Fab nanobody (*29*) was synthesized by GeneArt (Thermo Fisher Scientific) and subsequently cloned into the pET28a vector downstream of a 6xHis tag. The resulting fusion proteins were expressed in *Escherichia coli* BL21 (DE3) at 18°C overnight. Protein expression was induced with 0.5 mM Isopropyl ß-D-1-thiogalactopyranoside (IPTG) at an OD A_600_ of 0.6. The fusion proteins were first purified in batch by a Ni^2+^-affinity step (HisPur Ni-NTA Resin; Thermo Fisher Scientific), before being further purified by size-exclusion chromatography (Superdex 75 Increase 10/300 GL). The final protein buffer solution contained 25 mM Tris-HCl (pH 7.5) and 150 mM NaCl (no reducing agents). Monomeric anti-Fab nanobody fractions were pooled, concentrated to ∼3.8 mg/ml, flash-frozen in liquid nitrogen, and stored at -70°C.

### Purification of the NabFab

The NabFab plasmid was kindly provided by the group of Prof. Kaspar Locher, ETH Zürich (*29*). Chemically competent C43 *Escherichia coli* cells were used for protein expression. Four liters of TB autoinduction media (Terrific Broth containing 0.4% glycerol, 0.01% glucose, 0.02% lactose,1.25 mM MgSO_4_ and 100 μg/ml ampicillin) were inoculated with overnight cultures from single colonies and incubated for 6 h at 37 °C, before shifting to 30°C for expression overnight at 180 rpm shaking.

Cells were resuspended in lysis buffer containing 50 mM Tris-HCl pH 7.5, 200 mM NaCl, protease inhibitor cocktail tablets (PIC) (complete EDTA-free, Roche Diagnostics) and 5 units/ml supernuclease (Novagen) and lysed by sonication. The lysate was incubated in a water bath at 65°C for 40 min and spinned down at 20,000x g for 30 min. The filtered supernatant was loaded on a HiTrap™ Protein L column (Cytiva), pre-equilibrated with 50 mM Tris-HCl (pH 7.5) and 500 mM NaCl. The protein was eluted using 0.1 M acetic acid and immediately loaded onto a Hitrap SP HP (Cytiva) column pre-equilibrated with buffer A (50 mM sodium acetate, pH 5.0). After a washing step, the NabFab was eluted using a salt gradient with buffer B (50 mM sodium acetate, 2 M NaCl, pH 5.0). The final protein buffer contained 25 mM Tris-HCl pH 7.5, 150 mM NaCl, the protein was concentrated to 7.0 mg/ml, flash-frozen and stored at -70°C.

### Purification of µOR

DNA coding for the full-length murine µOR was subcloned into the pF1 vector. The resulting construct comprises an N-terminal signal sequence, a Flag epitope tag and a C-terminal HRV-3C protease cleavage site followed by a Strep II tag and 8 x His tag. Sf9 cells were used for expression of µOR in the presence of 10 µM naloxone. Cells were infected at a cell density of roughly 3.0 × 10^6^ cells/ml, incubated for 48 h at 27°C at 110 rev/min and harvested by centrifugation at a cell viability of 80–85%. The purification of µOR has been previously described (*25*). All purification steps were performed at 4°C. Briefly, Sf9 cell pellets were lysed using hypotonic buffer containing 20 mM HEPES pH 7.5, 5 mM MgCl_2_, PIC, 5 units/ml supernuclease, 10 μM naloxone (Sigma-Aldrich) and 2 mg/ml iodoacetamide (Sigma-Aldrich), and gently stirred for 1 h. Cell membranes were separated by ultracentrifugation at 100,000x g for 40 minutes and resuspended in solubilization buffer made of 25 mM HEPES pH 7.5, 500 mM NaCl, 1% LMNG/0.1% CHS, PIC, 10 μM naloxone and 2 mg/ml iodoacetamide using a dounce homogenizer. The solution was gently stirred for 4 h. The insoluble debris was removed by ultracentrifugation at 100,000x g for 1 h. The supernatant (supplemented with 20 mM imidazole) was incubated for 2-3 hours with HisPur Ni-NTA Resin and extensively washed with wash buffer A (25 mM HEPES pH 7.5, 500 mM NaCl, 0.1% LMNG/0.01% CHS, 1 μM naloxone, 10 mM imidazole), followed by buffer B (25 mM HEPES pH 7.5, 500 mM NaCl, 0.1% LMNG/0.01% CHS, 1 μM naloxone, 10 mM MgCl_2_, 10 mM ATP). Proteins were eluted using an imidazole gradient and subsequently loaded onto a 5 ml StrepTactin Superflow Cartridge (Qiagen) at a flow rate of 0.8 ml/min. The column was washed with wash buffer C (25 mM HEPES pH 7.5, 500 mM NaCl, 0.05% LMNG/0.005% CHS) to remove contaminations and residual naloxone. µOR was eluted using 2.5 mM desthiobiotin. The µOR was further purified by size exclusion chromatography. A Superose 6 Increase 10/300 GL column (GE Healthcare Life Sciences) was pre-equilibrated with 20 mM HEPES pH 7.5, 100 mM NaCl, and 0.001% LMNG/0.0001% CHS. Fractions containing µOR were concentrated, flash-frozen and stored at -80°C.

### NbE-µOR-Fab module complex assembly

To assemble the NbE-µOR-Fab module complex, freshly prepared µOR was incubated with NbE, NabFab and anti-Fab Nb in a molar ratio of 1:2:2.5:3 overnight, gently mixed at 4°C. A final size-exclusion step using a Superose 6 Increase 10/300 GL column was conducted. Fractions containing the complex were pooled, concentrated to around 2.8 mg/ml and immediately used for EM grid preparation.

### Cryo-EM sample preparation and data collection

3 µl of freshly purified NbE-µOR-Fab module complex were applied onto holey gold grids (Quantifoil UltrAuFoil R1.2/1.3, 300 mesh), front blotted for 3-4 seconds with 1 mm additional movement (95% humidity at 15°C) before being plunged into liquid ethane using an EM GP2 automatic plunge freezer (Leica). The cryo-EM data sets of NbE-µOR-Fab module complex were acquired on a Thermo Scientific Talos Arctica Cryo-TEM at an accelerating voltage of 200 kV. A total of 5’938 movies were recorded using a Falcon III direct electron detector at a nominal magnification of 150,000x, resulting in a pixel size of 0.9759 Å. Data were collected using EPU (Thermo Fisher Scientific) with one image per hole, a defocus range of −0.6 to −2.0 μm and a total electron dose of 40 e^−^/Å^2^ distributed over 44 frames per acquisition. Data acquisition was monitored on-the-fly pre-processing software using CryoSPARC v.3.3.1(*43*)

### Cryo-EM image processing

All data were processed using CryoSPARC v.3.3.1 and RELION 3.1(*43*, *46*). The data processing workflow is summarized in Suppl. Figure S3. First, raw movies were aligned and dose weighted using patch-based motion correction (*47*). Contrast transfer function (CTF) parameters were estimated by patch-based CTF estimation (*43*). Only micrographs with a CTF fit better than 4.0 Å resolution were selected for further processing, resulting in a set of 5’686 micrographs. Particles were initially picked using a blob picker with a minimum and maximum diameter of 70 Å and 200 Å, respectively. The resulting 2D classes were fed into the Topaz particle-picking pipeline to increase the number and accuracy of picked particles. After several rounds of 2D classification (and removal of duplicated particles) 704’786 particles were selected. *Ab initio* reconstruction with multiple classes was followed by heterogeneous refinement. The best class contained 509,809 particles. 3D classification helped to further clean up the particle set. 3D refinement using non-uniform refinement and CTF refinement (global and local CTF refinement) resulted in a reconstruction of 3.1 Å resolution. To further improve the density of the µOR, refined particles were imported into RELION 3.1 for 3D classification without alignment (K=4 and T=12), using a soft mask on the seven transmembrane helices of the µOR. The best class showing high resolution features was selected (445,766 particles) and particles were re-imported to CryoSPARC for non-uniform refinement followed by local refinement with a soft mask around the NbE-µOR. As a result, the density for the NbE-µOR interface was significantly improved. All maps were sharpened with deepEMhancer (*44*). All resolution estimations were derived from Fourier shell correlation (FSC) calculations between reconstructions from two independently refined half-sets, and reported resolutions are based on the FSC = 0.143 criterion. Local resolution estimations are obtained by ResMap (*48*).

### Cryo-EM model building and refinement

The crystal structure of inactive µOR bound to a covalent antagonist (PDB: 4DKL) was used as an initial reference (*25*). Similarly, the crystal structure of the NbE published in this manuscript was used as an initial model. For NabFab and anti-Fab Nb, the cryo-EM structure of VcNorM complex was used as initial models (*29*). All initial models were fitted into cryo-EM maps using Chimera X (*49*), then manually built in Coot (*50*), iteratively and real-space refined using PHENIX (*51*). Model validation was performed with MolProbity (*52*). Structural figures were generated in Chimera X (*49*).

### Crystallization and data collection

Purified NbE was concentrated to ∼30 mg/mL for crystallization. Crystals were grown using the hanging drop vapor diffusion method at 16°C in a temperature-controlled incubator. The best NbE crystals grew in 1 M succinic acid pH 7.0, 1-2% polyethylene glycol 2000 monomethyl ether (PEG 2000 MME) and 0.1 M Hepes, pH 7.0, reaching their final size within 3 days. Crystals of NbE were cryoprotected using the well solution supplemented with 25% glycerol and flash frozen in liquid nitrogen. Diffraction data was collected at the Advanced Photon Source on beamline 23-IDD on a Pilatus 6M detector. The final dataset was collected from a single crystal of NbE, with reflections extending to 2.85 Å resolution.

### Crystallographic data processing and model refinement

Reflection data was integrated using XDS (*53*), and scaled and merged using Aimless as part of the CCP4 suite (*54*). Initial phases were obtained using molecular replacement in Phaser (*55*), using a homology model from SwissModel (*56*) yielding three copies of NbE in the asymmetric unit. Iterative rounds of manual model building and automated refinement were carried out in Coot (*57*) and Phenix (*51*), respectively. The final model was refined to a R_free_ of 0.331 with favorable geometry (96% Ramachandran favored, 3% allowed, and 0.3% outliers). Of the three chains in the asymmetric unit, one (chain A) is characterized by exemplary density for the resolution and was used as an initial model for building of the NbE-µOR-Fab complex.

### Peptide synthesis

Peptides have been synthesized using Fmoc-based solid phase peptide synthesis (SPPS) on a microwave assisted peptide synthesizer (CEM Liberty Lite). The synthesis was performed on 0.1 mmol scale using preloaded Wang resin or Rink Amide resin depending on the desired C-terminal end of the peptide, being a carboxylic acid or carboxamide. Fmoc deprotection was performed at 90°C for 3 min using a solution of 20% 4-methylpiperidine in DMF during the entire synthesis. Each coupling was done using 5 equivalents of Fmoc protected amino acid, with 0.5 M DIC and 1 M Oxyma as coupling reagents. N-terminal acetylation has been done manually using 10 equivalents of acetic anhydride and 5 equivalents of DIPEA during 1 h in DMF. After completion of the sequence, the resin was washed several times with DCM, followed by the cleavage using a cocktail solution consisting of 90 % TFA, 5 % triisopropylsilane and 5 % distilled water during 4 h. After freeze-drying, crude peptides were obtained and purified using preparative HPLC. More specifically, a Gilson HPLC system, equipped with Gilson 322 pumps and a Supelco Discovery® BIO Wide Pore C18 column (25 cm x 21.2 mm, 10 µm), was used. Crude peptides were dissolved in DMSO and purified using H_2_O-AcN–0.1% TFA as mobile phase. Finally, fractions were collected, and the accompanying purities were assessed by analytical RP-HPLC, after which the pure fractions were combined and lyophilized to obtain the final purified peptide as a powder (TFA salt) with a high purity (> 95 %).

### Data availability

The EM maps of the NbE-µOR-Fab module complex have been deposited in the Electron Microscopy Data Bank (EMDB) under the accession code EMD-18541. Protein coordinates for NbE-µOR-Fab module complex structure and the crystal structure of the isolated nanobody NbE have been deposited in the Protein Data Bank (PDB) under accession codes 8QOT and 8V8K, respectively. Source data are provided with this paper.

## Acknowledgements

We thank A. Radoux for valuable advice and discussion, and M. de Lapeyrière for generating preliminary data on NbE’s antagonism. We thank the Bioimaging, Flow Cytometry, PPR2P and the RE.A.D.S Core Facilities at the University of Geneva. We thank Y. Pfister for technical assistance, A. Howe for assistance with EM data collection at the DCI-Geneva (cryoGEnic) EM facility, the computing department for providing the infrastructure to perform cryo-EM analysis, N. Roggli for maintaining computing in the Molecular and Cellular Biology department, and C. Bauer for his contributions to the cryoGEnic facility. This work was supported by the Swiss National Science Foundation (Eccellenza Professorial Fellowship PCEFP3_181282 to M.S. and research grants 310030_185235 and TMSGI3_211581 to A.B.), a grant of the Helmut Horten foundation, and a grant to M.S. and A.B. of the Boninchi Foundation. C.M. and S.B. acknowledge the Research Council of the Vrije Universiteit Brussel for the financial support through the Strategic Research Program (SRP50). We thank members of the Stoeber and Boland group for critical comments on the manuscript.

Molecular graphics and analyses performed with UCSF ChimeraX, developed by the Resource for Biocomputing, Visualization, and Informatics at the University of California, San Francisco, with support from National Institutes of Health R01-GM129325 and the Office of Cyber Infrastructure and Computational Biology, National Institute of Allergy and Infectious Diseases.

## Author contributions

J.Y. purified and reconstituted the NbE-µOR-NabFab complex and determined the cryo-EM structure. A.K. performed the functional characterizations of NbE, peptidomimetics, and receptor mutants. X.Z and P.R. performed GCI affinity measurements. C.M. and S.B designed and synthesized peptide mimetics. A.M. and A.K. provided NbE and solved its crystal structure in isolation. All authors planned experiments and analyzed data. T.L. and J.S. generated NbE and characterized the binding. A.B. and M.S. supervised the project and wrote the manuscript with input from all other authors.

## Competing interests

The authors declare no competing interests.

---

**Supplementary figure S1:**
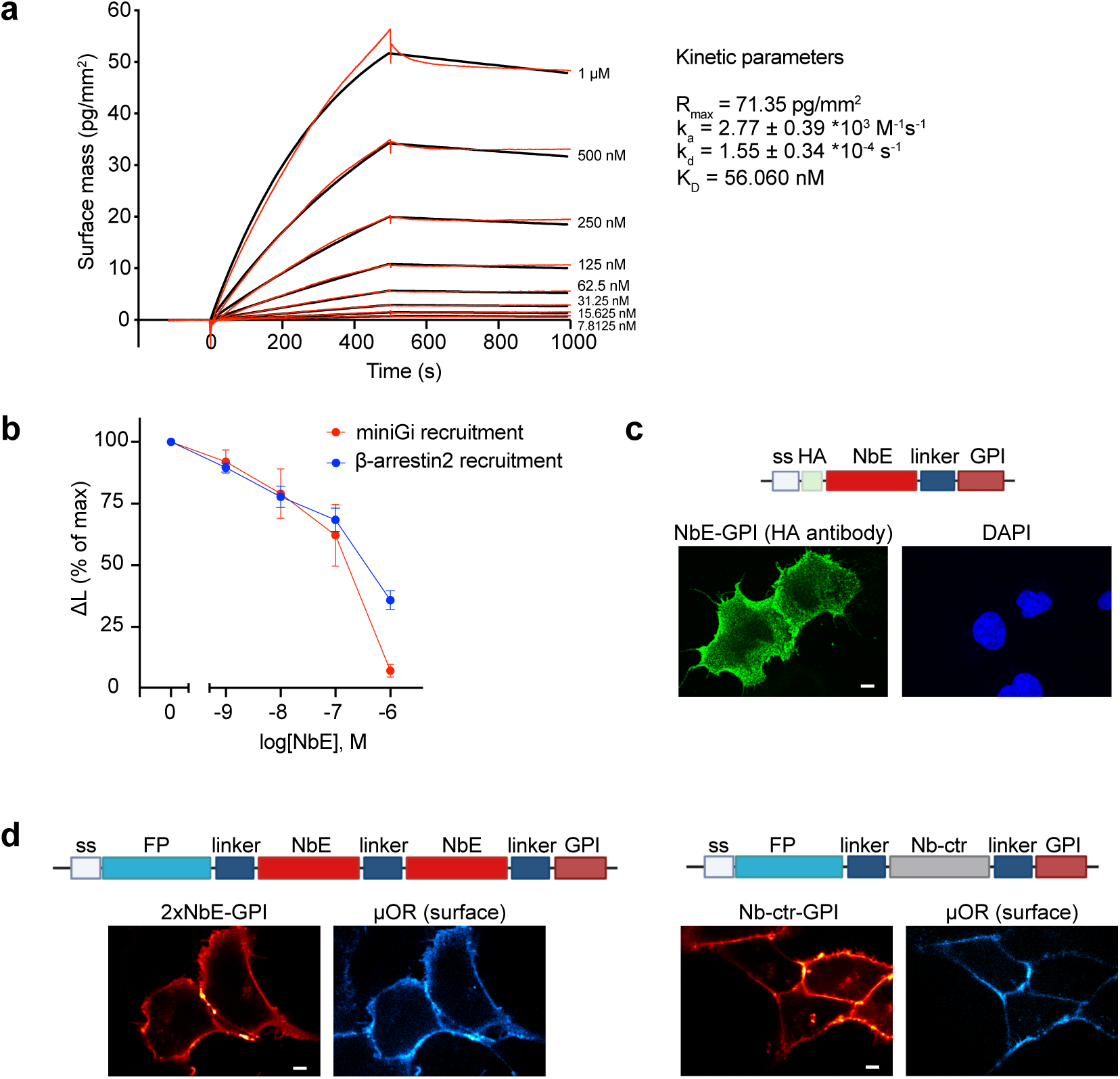
Functional characterization of NbE. **(a)** Grating coupled interferometry (CGI)-based kinetic data of μOR binding to AVI-tagged NbE, immobilized on a Creoptix WAVEchip. Measurements obtained for μOR concentrations between 7.8 nM to 1000 nM. Rmax is the maximal response value, ka and kd are the association and dissociation rate constants, KD is the dissociation constant. **(b)** DAMGO-driven recruitment of mGi1-LgBiT (100 nM DAMGO) or β-arrestin2-LgBiT (1 μM DAMGO) to μOR-SmBiT (NanoLuc complementation assay). Cells were pre-incubated with different concentrations of NbE. N ≥ 3, mean ± SEM. **(c)** Confocal images of HEK293 cells transfected with HA-NbE-GPI and stained with AF488-conjugated anti-HA antibody in non-permeabilizing conditions (green). DAPI staining is shown (blue). Scale bar, 10 μm. **(d)** Confocal images of HEK293 cells stably expressing μOR (labeled with anti-FLAG M1-AF647, cyan) and transfected with mRuby2-2xNbE-GPI (left) or a mRuby2 control Nb-GPI (right) (red). Scale bars, 10 μm.

**Supplementary figure S2:**
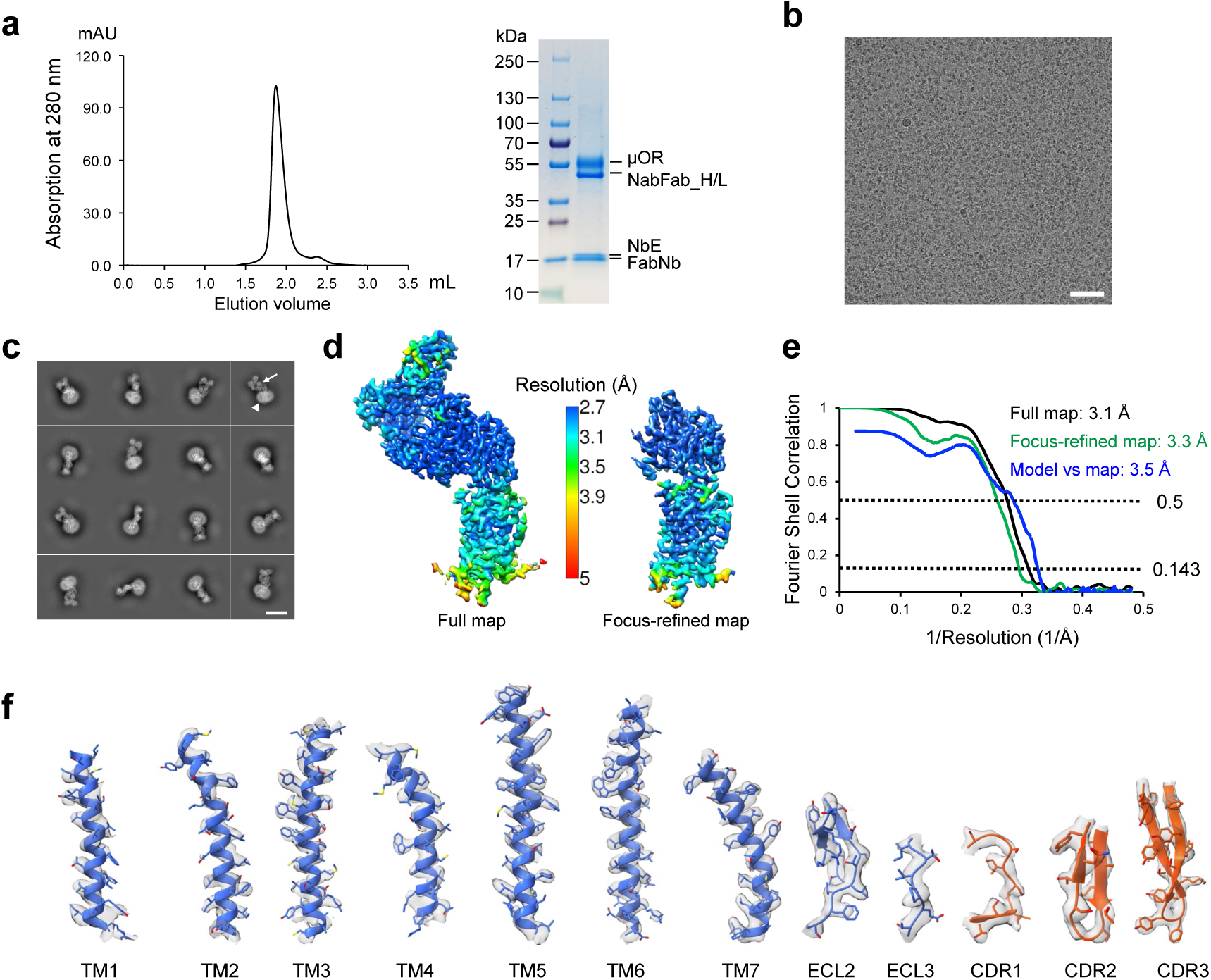
Structure determination of the NbE-µOR-Fab module complex. **(a)** Size exclusion chromatogram (left) and SDS–PAGE gel (right) of the NbE-µOR-Fab module complex, demonstrating the formation of a stable complex. **(b)** Representative cryo-EM micrographs of the NbE- µOR-Fab module complex collected on UltrAuFoil EM grids. Scale bar, 50 nm. **(c)** Gallery of two-dimensional class averages of the NbE-µOR-Fab module complex showing typical 2D classes. White arrow, Fab module complex. White triangle, µOR in a LMNG micelle. Scale bar, 100 Å. **(d)** EM density maps color-coded according to local resolution, with the full map depicted on the left and the NbE-µOR focus refined map shown on the right. **(e)** Gold standard Fourier Shell Correlation (FSC) curves of the full map and the focused refined map (focussing on the NbE-µOR complex) and FSC curve between the full cryoEM map and the final atomic coordinates of the NbE-µOR-Fab module complex calculated using Mtriage as part of PHENIX package (*42*). **(f)** Representative EM density for regions of the µOR and NbE. Displayed are densities of all transmembrane helices, parts of ECL2 and ECL3 of the µOR and parts of CDR1, CDR2 and CDR3 of NbE, with the respective coordinates fitted.

**Supplementary figure S3:**
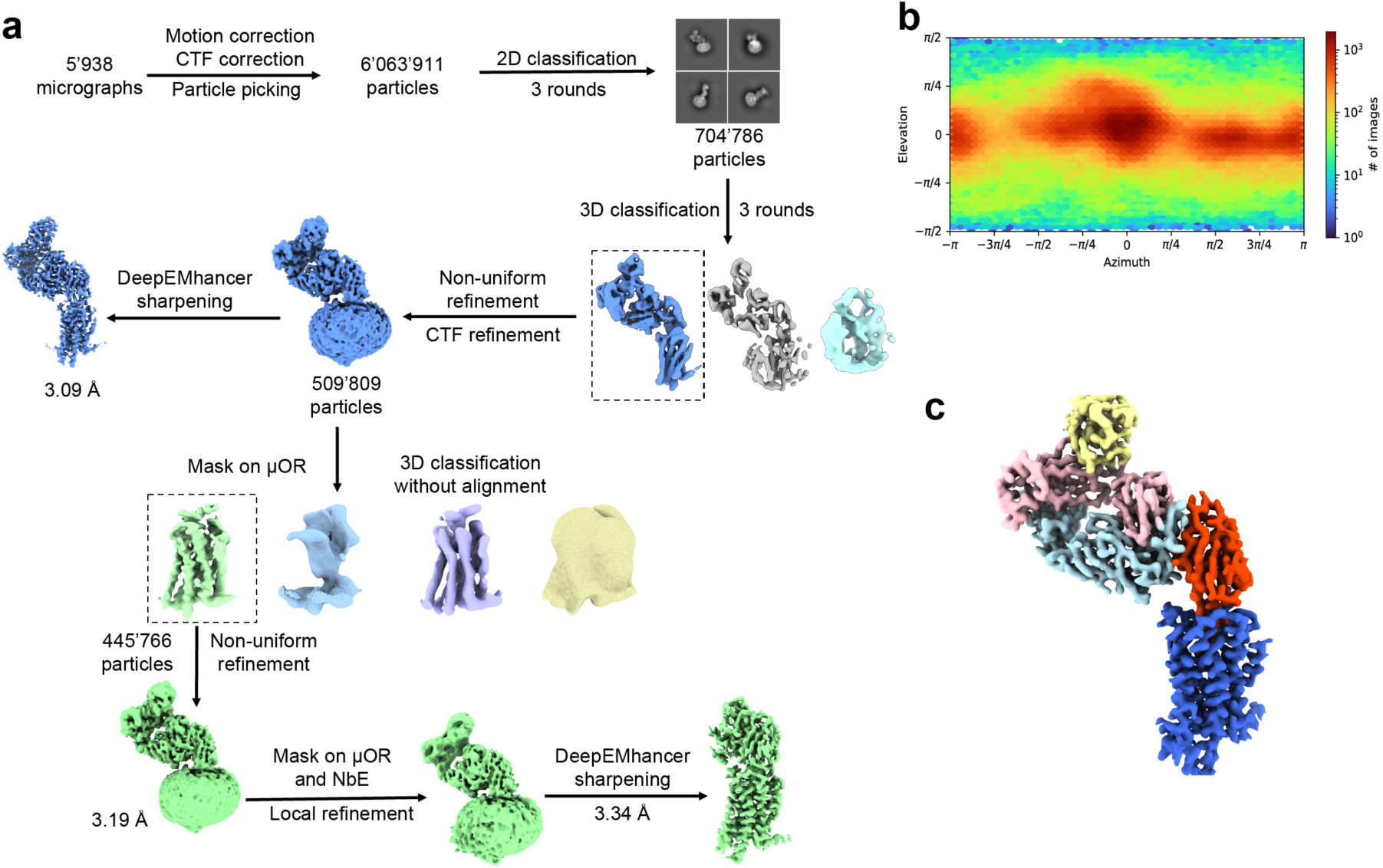
Data-processing flowchart for the NbE-µOR-Fab module complex. **(a)** See Methods: Roughly six million particles were picked from almost 6’000 EM micrographs. Data clean-up by 2D classification resulted in 704’786 particles that were further classified by three rounds of 3D classification. These 509’809 particles were subjected to CTF refinement and a mask for 3D classification was applied on the µOR. This resulted in a final set of 445’766 particles that was refined using a mask on the NbE-µOR complex, to improve the interface density. The final map was refined to 3.3 Å. **(b)** Angular distribution plot calculated in CryoSPARC (*43*). **(c)** Composite map of the NbE-µOR- Fab module complex sharpened with DeepEMhancer (*44*).

**Supplementary figure S4:**
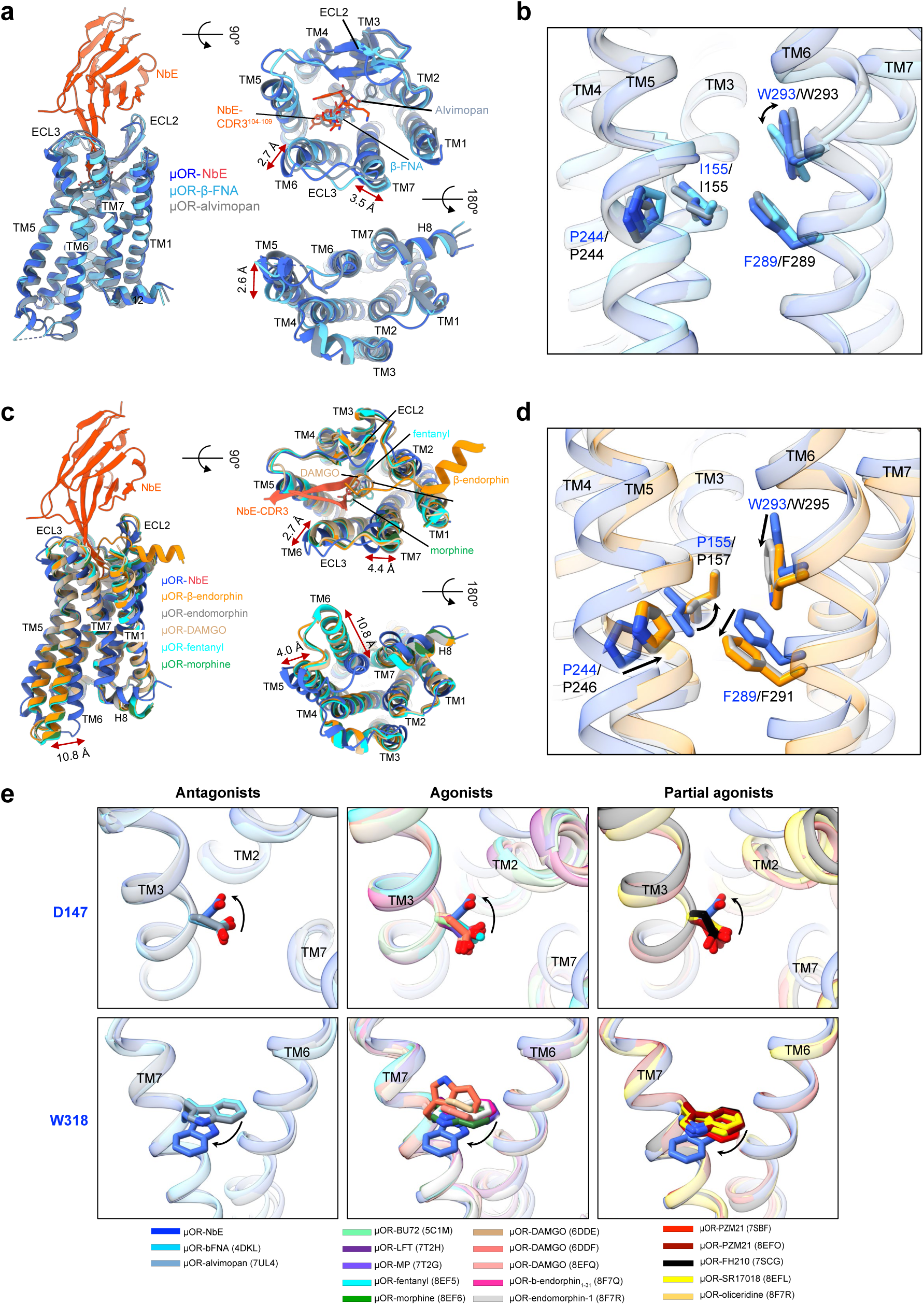
Conformational changes of the µOR induced by NbE ligand binding. **(a)** Superpositions of the µOR bound to the antagonists NbE, β-FNA (PDB: 4DKL) or alvimopan (PDB: 7UL4). The overall architecture superimposes well between the antagonists-bound structures (left). Binding of NbE leads to noticeable shifts in TM6, ECL3, TM7 near the orthosteric binding pocket (top view, right), and the intracellular tip of TM5 (bottom view, right). **(b)** The conserved core triad F289^6.44^, P244^5.50^ and I155^3.40^, superimposes well between the three antagonist-bound structures. **(c)** Superpositions of the µOR bound to NbE or the agonistic peptides β-endorphin (PDB: 8F7Q) endomorphin (8F7R), and DAMGO (6DDE), or small molecules morphine (8EF6) and fentanyl (8EF5). The intracellular part of TM6 is markedly shifted between the active (agonist-bound) form and the inactive (NbE-bound) form (bottom view, right). Binding of NbE leads to noticeable shifts in TM6, ECL3, TM7 near the orthosteric binding pocket (top view, right), and the intracellular tip of TM5 (bottom view, right). **(d)** The conserved core triad F289^6.44^, P244^5.50^ and I155^3.40^, is rearranged in the active conformation of the receptor. **(e)** Superposition of the rotamer conformations of D147^3.32^ and W318^7.35^ in all experimentally determined µOR structures. D147^3.32^ shows a unique rotamer conformation when bound to NbE compared to antagonists-, agonists- or partial agonists-bound structures (top panel). W318^7.35^ also adopts a unique position when bound to NbE (rare rotamer conformation and shifted, lower panel).

**Supplementary figure S5:**
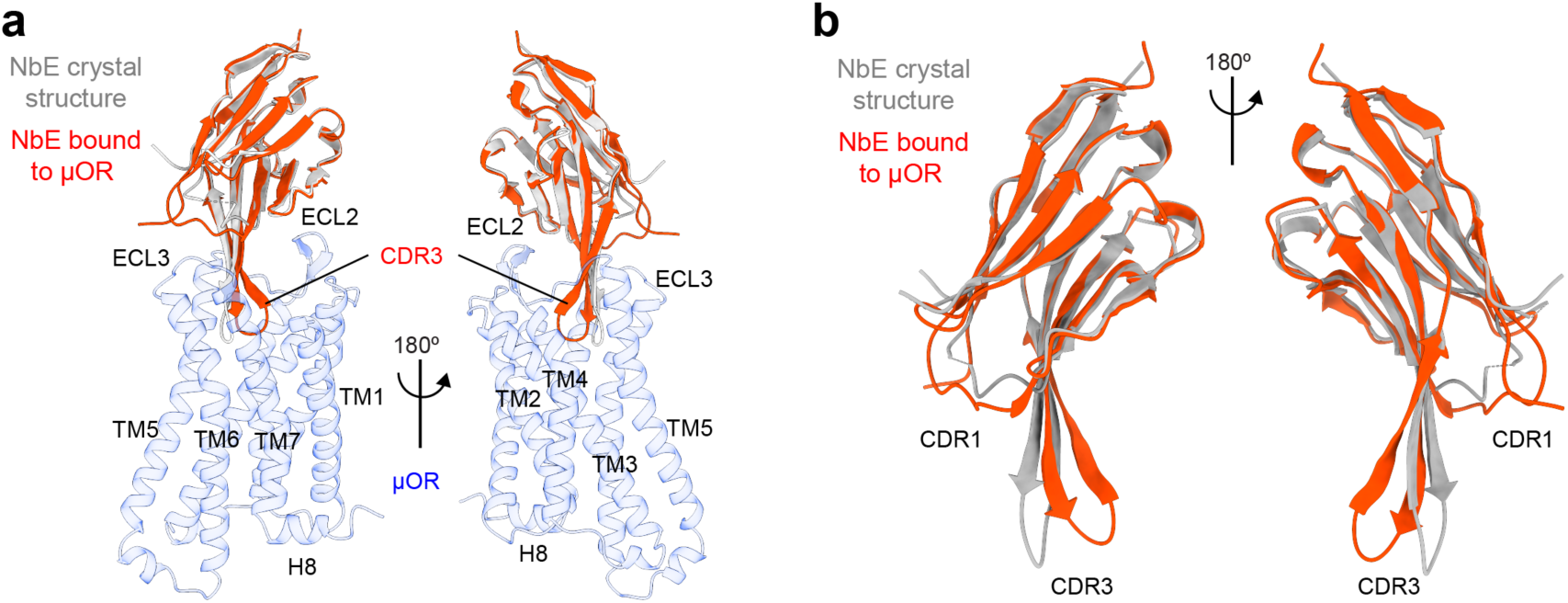
Structural comparison between NbE bound to the µOR or crystallized in isolation. **(a)** The NbE crystal structure is superimposed on the NbE-µOR complex structure, with the NbE bound to the µOR shown in red, the crystallized NbE form in gray and the µOR in light blue. The CDR3 conformation observed in the crystal structure would clash with the orthosteric binding pocket. **(b)** Superposition of the conformations of NbE, bound to the µOR (red) or in isolation (gray).

**Supplementary figure S6:**
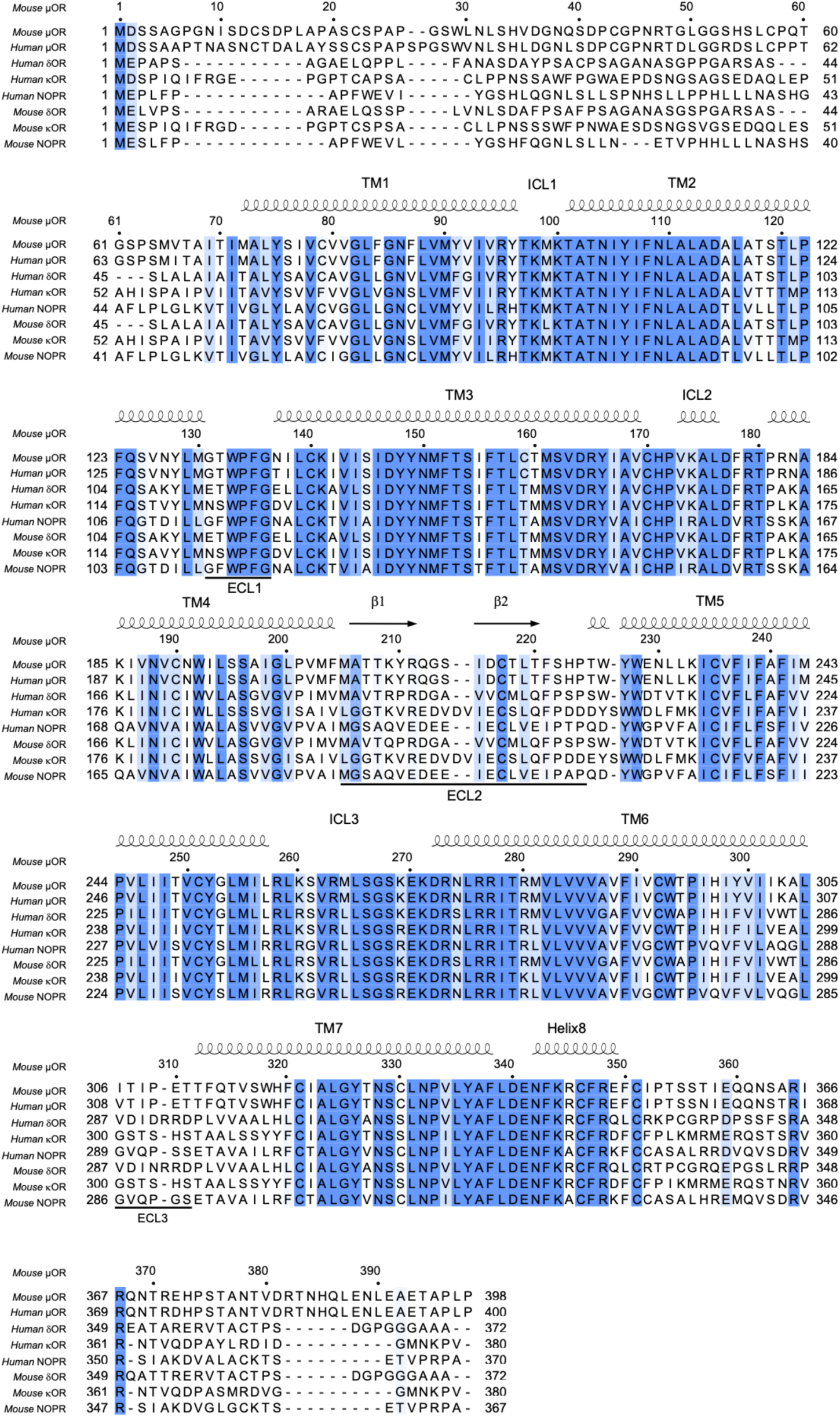
Multiple sequence alignments (MSA) of all four OR family members in mouse and human. Sequence alignment of the mouse µOR with all four family members of human OR, and mouse δOR, κOR, or NOPR. Degree of sequence conservation is color-coded according to ClustalX, and secondary structure elements are indicated above the sequences using ESPript 3 (*45*). ECL2 and ECL3 are indicated below the MSA. UniProt accession numbers: µOR (human P35372, murine P42866). δOR (human P41143, murine P42867), κOR (human P41145, murine P32300), NOPR (human P41146, murine P35377).

**Supplementary figure S7:**
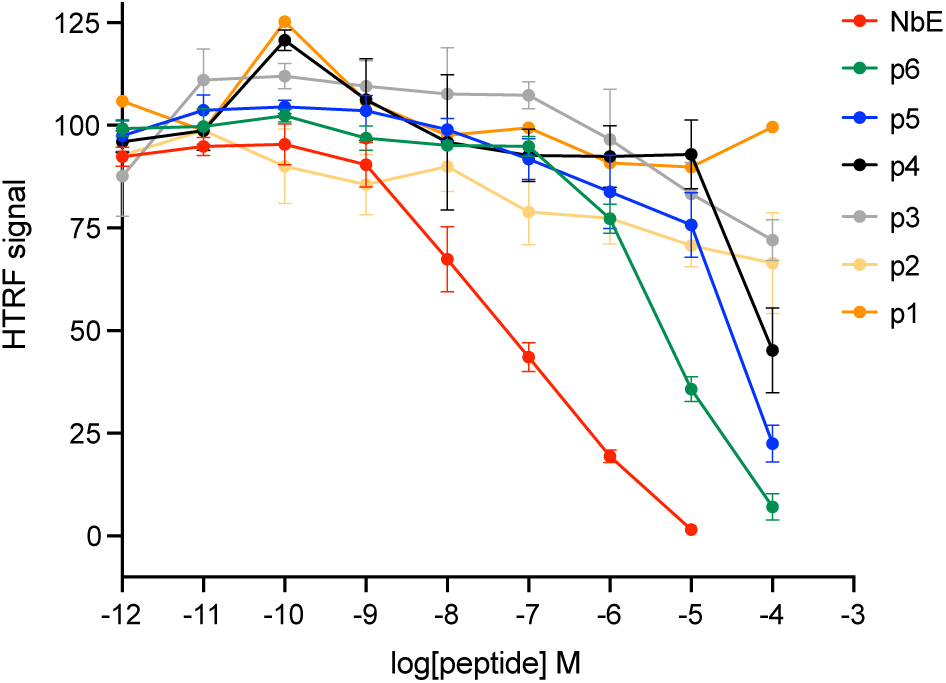
HTRF competition binding assay profiles of peptides (#1 - #6). HEK293 cells stably expressing SNAP-μOR were labeled with SNAP-Lumi4-Tb and 3 nM red-labeled naltrexone derivative, and incubated with increasing concentrations of peptides. Normalized to cells without peptides; N ≥ 3, mean ± SEM.

**Supplementary figure S8:**
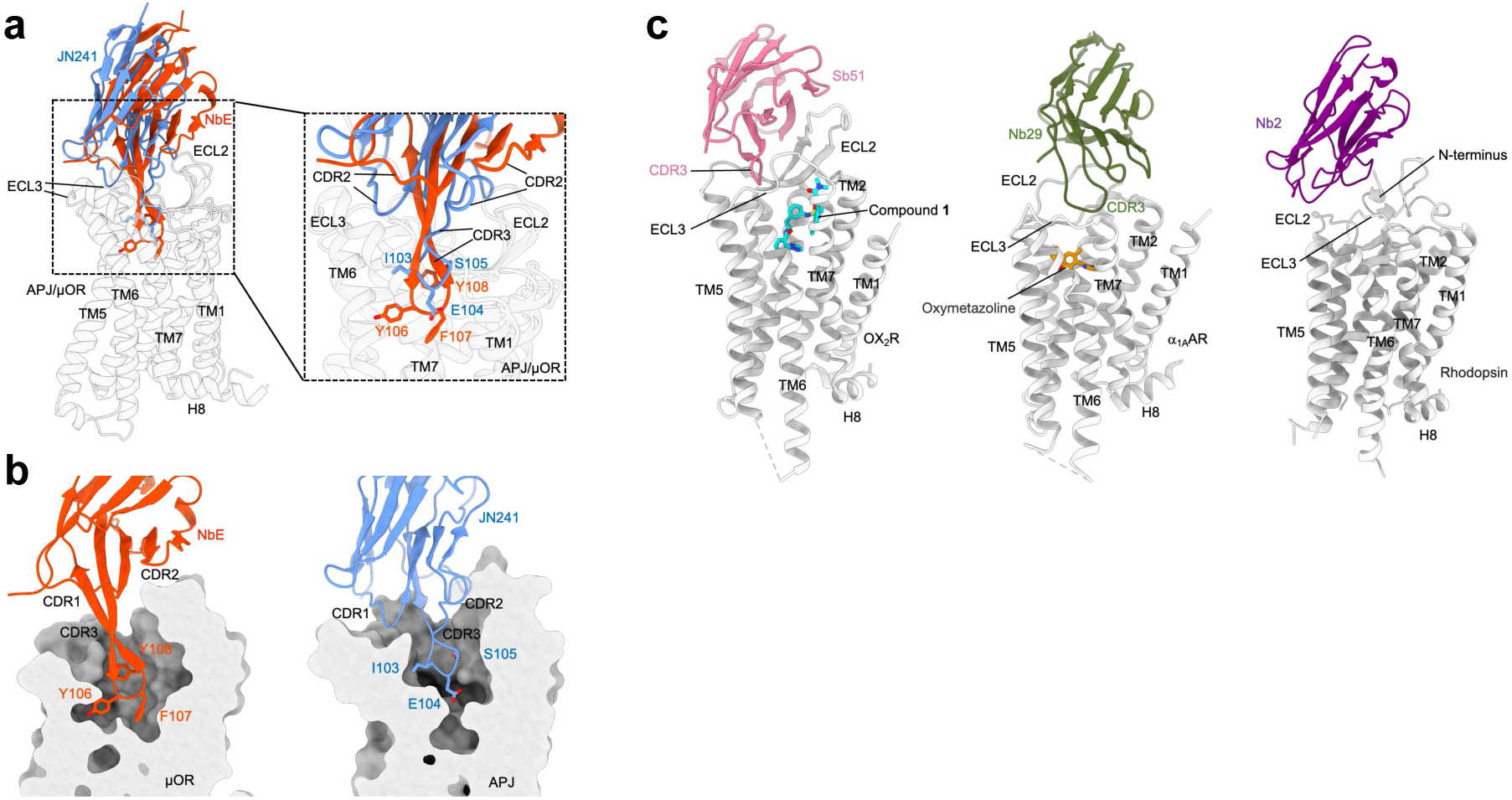
Comparison of extracellular nanobody binding modes to GPCR targets. **(a)** Superposition of Nbs JN241 (blue) with NbE (red). For reasons of clarity the human apelin receptor has been removed and only the µOR (opaque) is shown as a GPCR reference. **(b)** Binding of the CDR3s of NbE and JN241 to the µOR or the apelin receptor, respectively. Both CDR3s are deeply inserted into the ligand binding pocket, however, in contrast to NbE, the CDR3 of JN241 is not formed of aromatic residues important for receptor recognition. **(c)** Nbs Sb51 (pale violet) and Nb29 (olive green) bind to the solvent-exposed extracellular loops of the orexin receptor 2 (*37*) and the α1A-adrenergic receptor (*36*), respectively. Sb51 and Nb29 are positioned above the small molecule ligands compound 1 (cyan) and oxymetazoline (orange), which are bound to the receptors’ orthosteric pockets. Nb2 (purple) binds to the ECL2 and N-terminus of rhodopsin (*18*).

**Supplementary table 1:**
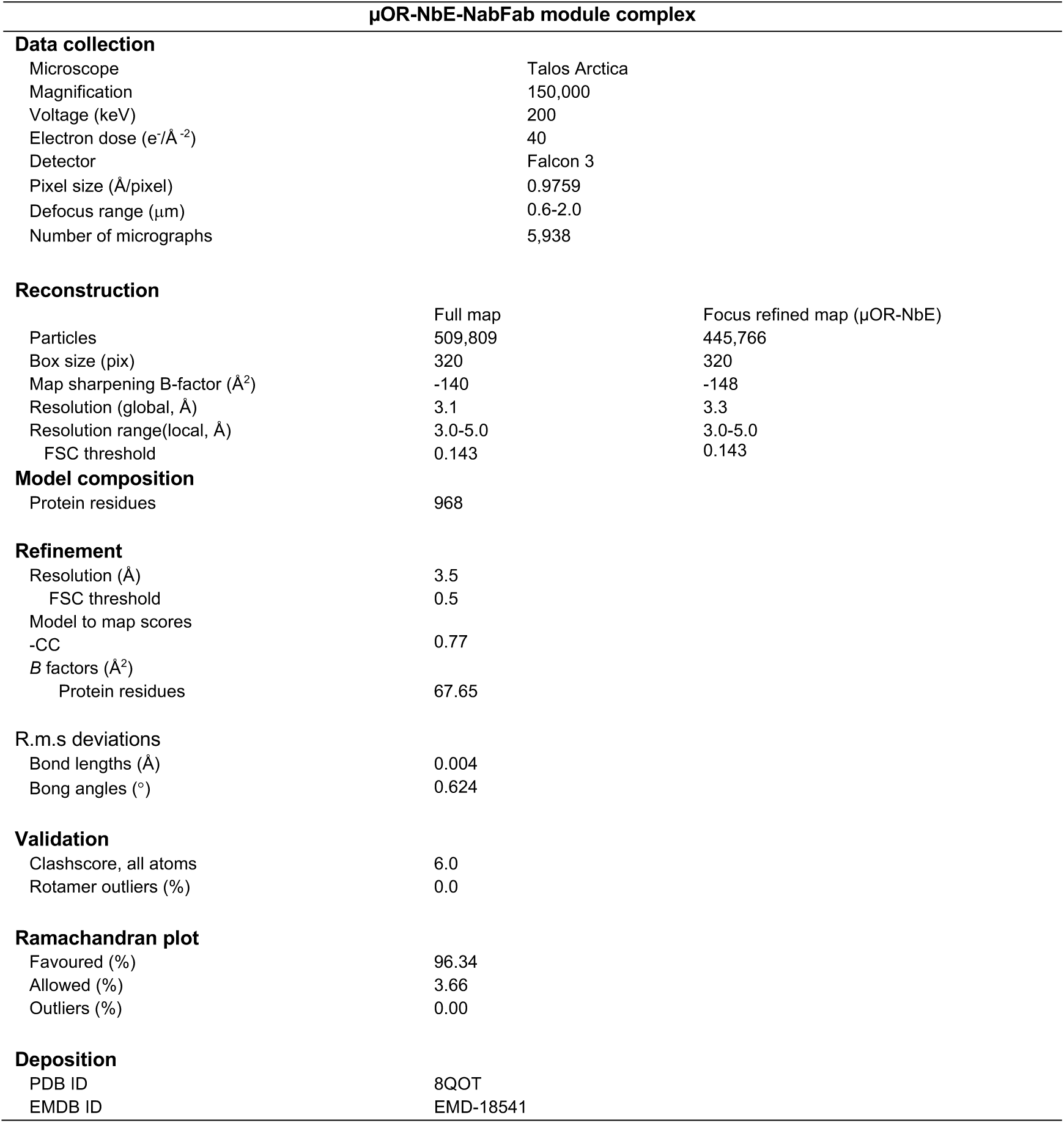
Cryo-EM data collection, refinement, and validation statistics.

**Supplementary table 2:** Information on structures of µOR bound to antagonists, full agonists, or partial agonists. The computed average root mean square deviation (RMSD) for all Cα atoms of µOR between the NbE-µOR complex and the respective µOR-ligand complexes is indicated.

| Ligand | Ligand type | RMSD C<sub>α</sub> atoms<br>(vs NbE-μOR complex) | Resolution<br>(in Å) | PDB ID |
| --- | --- | --- | --- | --- |
| *β*-FNA | Antagonist | 0.815 | 2.8 | 4DKL |
| *Alvimopan* | Antagonist | 0.749 | 2.8 | 7UL4 |
| *BU72* | Full agonist | 1.714 | 2.1 | 5C1M |
| *DAMGO* | Full agonist | 1.937 | 3.5 | 6DDE |
| *DAMGO* | Full agonist | 1.901 | 3.5 | 6DDF |
| *Mitragynine*<br>*pseudoindoxyl* | Full agonist | 1.739 | 3.2 | 7T2G |
| *Lofentanil* | Full agonist | 1.672 | 2.5 | 7T2H |
| *DAMGO* | Full agonist | 1.842 | 3.3 | 8EFQ |
| *Fentanyl* | Full agonist | 1.666 | 3.3 | 8EF5 |
| *Morphine* | Full agonist | 1.552 | 3.2 | 8EF6 |
| *β*-endorphin<sup>1-31</sup> | Full agonist | 1.595 | 3.2 | 8F7Q |
| *endomorphin-1* | Full agonist | 1.739 | 3.3 | 8F7R |
| *PZM21* | Partial agonist | 1.632 | 2.9 | 7SBF |
| *FH210* | Partial agonist | 1.964 | 3.0 | 7SCG |
| *Oliceridine* | Partial agonist | 1.933 | 3.2 | 8EFB |
| *SR17018* | Partial agonist | 1.568 | 3.2 | 8EFL |
| *PZM21* | Partial agonist | 1.711 | 3.2 | 8EFO |

**Supplementary table 3:** Residues in µOR interacting with agonists or antagonists. Residues in pink rows interact with all the five ligands analyzed.

| Agonists |  | Antagonists |  |  |
| --- | --- | --- | --- | --- |
| Endomorphin<br>(human μOR) | Beta-endorphin<br>(human μOR) | NbE<br>(mouse μOR) | Alvimopan<br>(mouse μOR) | Beta-FNA<br>(mouse μOR) |
|  | M67 (TM1) |  |  |  |
|  | I71 (TM1) |  |  |  |
| Y77 (TM1) |  |  |  |  |
| Q126 (TM2) | Q126 (TM2) |  | Q124 (TM2) |  |
| N129 (TM2) | N129 (TM2) |  | N127 (TM2) |  |
| Y130 (TM2) |  |  | Y128 (TM2) |  |
|  | L131 (TM2) |  |  |  |
|  | M132 (TM2) |  |  |  |
| W135 (ECL1) | W135 (ECL1) |  | W133 (ECL1) |  |
| V145 (TM3) | V145 (TM3) |  | V143 (TM3) |  |
| I146 (TM3) | I146 (TM3) |  | I144 (TM3) |  |
| D149 (TM3) | D149 (TM3) | D147 (TM3) | D147 (TM3) | D147 (TM3) |
| Y150 (TM3) | Y150 (TM3) | Y148 (TM3) | Y148 (TM3) | Y148 (TM3) |
| M153 (TM3) | M153 (TM3) | M151 (TM3) | M151 (TM3) | M151 (TM3) |
|  |  | K209 (ECL2) |  |  |
|  | R213 (ECL2) | R211 (ECL2) |  |  |
|  | Q214 (ECL2) | Q212 (ECL2) |  |  |
|  | D218 (ECL2) |  |  |  |
| C219 (ECL2) | C219 (ECL2) |  |  |  |
|  |  | L219 (ECL2) |  |  |
|  |  | F221 (ECL2) |  |  |
|  |  | E229 (TM5) |  | E229 (TM5) |
|  |  | L232 (TM5) |  |  |
| K235 (TM5) | K235 (TM5) | K233 (TM5) |  | K233 (TM5) |
| V238 (TM5) | V238 (TM5) | V236 (TM5) | V236 (TM5) | V236 (TM5) |
|  |  | W293 (TM6) |  | W293 (TM6) |
| I298 (TM6) | I298 (TM6) | I296 (TM6) | I296 (TM6) | I296 (TM6) |
| H299 (TM6) | H299 (TM6) | H297 (TM6) | H297 (TM6) | H297 (TM6) |
| V302 (TM6) | V302 (TM6) | V300 (TM6) | V300 (TM6) | V300 (TM6) |
|  |  | K303 (TM6) |  | K303 (TM6) |
|  | E312 (ECL3) | E310 (ECL3) |  |  |
| W320 (TM7) | W320 (TM7) | W318 (TM7) | W318 (TM7) | W318 (TM7) |
| H321 (TM7) |  |  |  |  |
| I324 (TM7) | I324 (TM7) | I322 (TM7) | I322 (TM7) | I322 (TM7) |
| Y328 (TM7) | Y328 (TM7) | Y326 (TM7) | Y326 (TM7) | Y326 (TM7) |

**Supplementary table 4:** Crystallographic data-collection and refinement statistics.

|  | NbE |
| --- | --- |
| **Data collection** |  |
| Space group | P6<sub>4</sub> 2 2 |
| Cell dimensions |  |
| *a*, *b*, *c* (Å) | 76.56, 76.56, 298.60 |
| $\alpha$, $\beta$, $\gamma$ (°) | 90, 90, 120 |
| Resolution (Å) | 44.37-2.85 (2.95-2.85) * |
| *R*<sub>meas</sub> | 0.175 (1.77) |
| CC<sub>1/2</sub> | 0.996 (0.536) |
| *I* / σ*I* | 12.3 (1.23) |
| Completeness (%) | 98.66 (95.31) |
| Redundancy | 11.9 (13.1) |
| **Refinement** |  |
| Resolution (Å) | 44.37-2.85 |
| No. reflections | 12826 (1198) |
| *R*<sub>work</sub> / *R*<sub>free</sub> | 0.286 / 0.331 |
| No. atoms |  |
| Protein | 2701 |
| Ligand/ion | 0 |
| Water | 0 |
| *B*-factors |  |
| Protein | 102.71 |
| Ligand/ion | - |
| Water | - |
| R.m.s. deviations |  |
| Bond lengths (Å) | 0.004 |
| Bond angles (°) | 0.97 |

\* Values in parentheses are for highest-resolution shell.

**Supplementary table 5:**
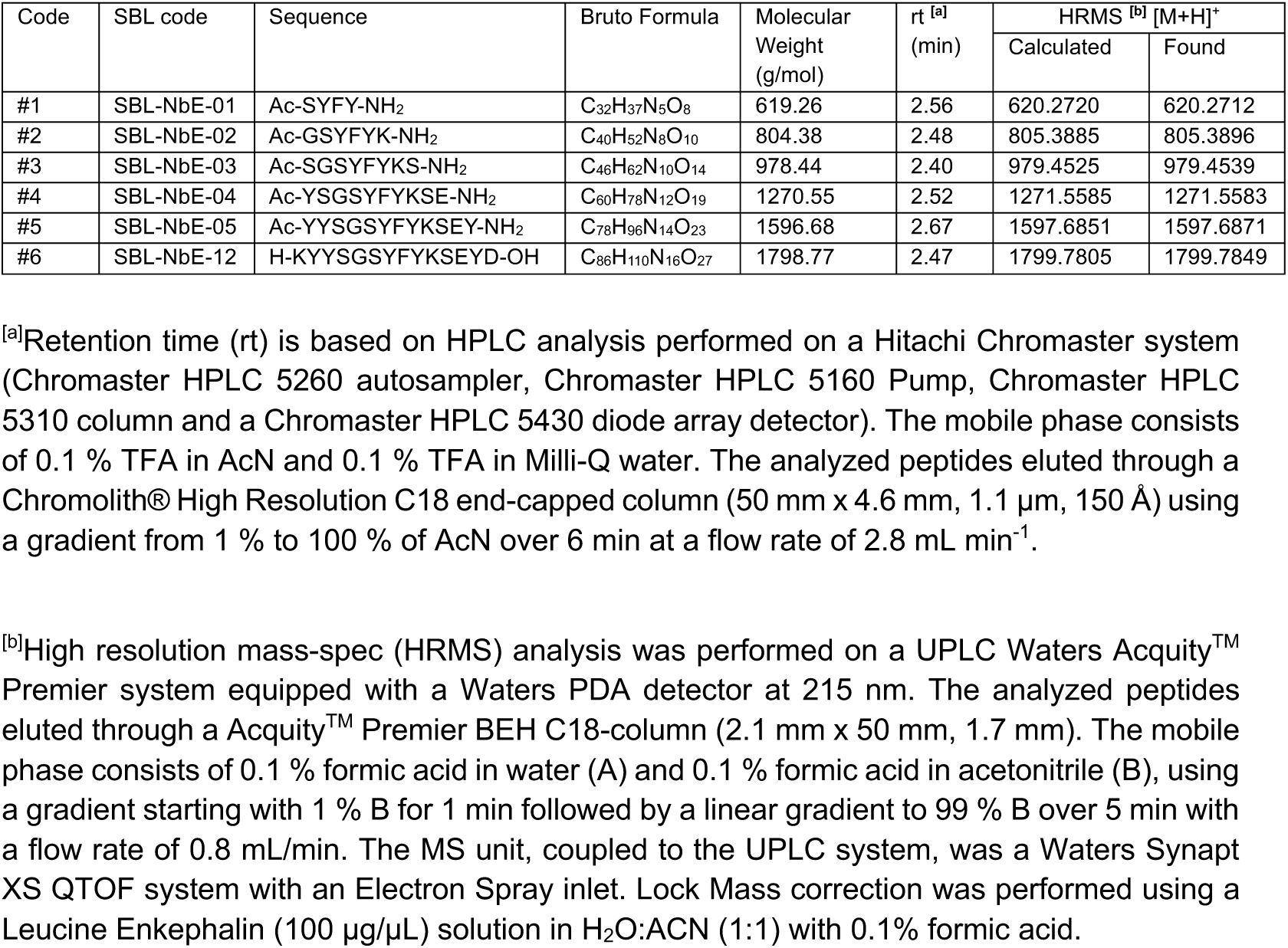
Analytical data of synthesized peptides.

## Notes

### Competing Interest Statement

The authors have declared no competing interest.

## References

1. R. Santos, O. Ursu, A. Gaulton, A. P. Bento, R. S. Donadi, C. G. Bologa, A. Karlsson, B. Al-Lazikani, A. Hersey, T. I. Oprea, J. P. Overington, A comprehensive map of molecular drug targets. Nat. Rev. Drug Discov. 16, 19–34 (2017).

2. K. Sriram, P. A. Insel, G Protein-Coupled Receptors as Targets for Approved Drugs: How Many Targets and How Many Drugs? Mol. Pharmacol. 93, 251–258 (2018).

3. A. S. Hauser, M. M. Attwood, M. Rask-Andersen, H. B. Schiöth, D. E. Gloriam, Trends in GPCR drug discovery: new agents, targets and indications. Nat. Rev. Drug Discov. 16, 829–842 (2017).

4. C. J. Hutchings, M. Koglin, W. C. Olson, F. H. Marshall, Opportunities for therapeutic antibodies directed at G-protein-coupled receptors. Nat. Rev. Drug Discov. 16, 661 (2017).

5. T. Laeremans, Z. A. Sands, P. Claes, A. De Blieck, S. De Cesco, S. Triest, A. Busch, D. Felix, A. Kumar, V.-P. Jaakola, C. Menet, Accelerating GPCR Drug Discovery With Conformation-Stabilizing VHHs. Front Mol Biosci. 9, 863099 (2022).

6. I. Jovčevska, S. Muyldermans, The Therapeutic Potential of Nanobodies. BioDrugs. 34, 11–26 (2020).

7. A. Manglik, B. K. Kobilka, J. Steyaert, Nanobodies to Study G Protein-Coupled Receptor Structure and Function. Annu. Rev. Pharmacol. Toxicol. 57, 19–37 (2017).

8. R. Heukers, T. W. M. De Groof, M. J. Smit, Nanobodies detecting and modulating GPCRs outside in and inside out. Curr. Opin. Cell Biol. 57, 115–122 (2019).

9. W. Huang, A. Manglik, A. J. Venkatakrishnan, T. Laeremans, E. N. Feinberg, A. L. Sanborn, H. E. Kato, K. E. Livingston, T. S. Thorsen, R. C. Kling, S. Granier, P. Gmeiner, S. M. Husbands, J. R. Traynor, W. I. Weis, J. Steyaert, R. O. Dror, B. K. Kobilka, Structural insights into µ-opioid receptor activation. Nature. 524, 315–321 (2015).

10. S. G. F. Rasmussen, H.-J. Choi, J. J. Fung, E. Pardon, P. Casarosa, P. S. Chae, B. T. Devree, D. M. Rosenbaum, F. S. Thian, T. S. Kobilka, A. Schnapp, I. Konetzki, R. K. Sunahara, S. H. Gellman, A. Pautsch, J. Steyaert, W. I. Weis, B. K. Kobilka, Structure of a nanobody-stabilized active state of the β(2) adrenoceptor. Nature. 469, 175–180 (2011).

11. M. J. Robertson, M. M. Papasergi-Scott, F. He, A. B. Seven, J. G. Meyerowitz, O. Panova, M. C. Peroto, T. Che, G. Skiniotis, Structure determination of inactive-state GPCRs with a universal nanobody. Nat. Struct. Mol. Biol. 29, 1188–1195 (2022).

12. A. C. Kruse, A. M. Ring, A. Manglik, J. Hu, K. Hu, K. Eitel, H. Hübner, E. Pardon, C. Valant, P. M. Sexton, A. Christopoulos, C. C. Felder, P. Gmeiner, J. Steyaert, W. I. Weis, K. C. Garcia, J. Wess, B. K. Kobilka, Activation and allosteric modulation of a muscarinic acetylcholine receptor. Nature. 504, 101–106 (2013).

13. R. Irannejad, J. C. Tomshine, J. R. Tomshine, M. Chevalier, J. P. Mahoney, J. Steyaert, S. G. F. Rasmussen, R. K. Sunahara, H. El-Samad, B. Huang, M. von Zastrow, Conformational biosensors reveal GPCR signalling from endosomes. Nature. 495, 534–538 (2013).

14. M. Stoeber, D. Jullié, B. T. Lobingier, T. Laeremans, J. Steyaert, P. W. Schiller, A. Manglik, M. von Zastrow, A Genetically Encoded Biosensor Reveals Location Bias of Opioid Drug Action. Neuron. 98, 963–976.e5 (2018).

15. C. McMahon, D. P. Staus, L. M. Wingler, J. Wang, M. A. Skiba, M. Elgeti, W. L. Hubbell, H. Rockman, A. C. Kruse, R. J. Lefkowitz, Synthetic nanobodies as angiotensin receptor blockers. Proceedings of the National Academy of Sciences. 117, 20284–20291 (2020).

16. P. Scholler, D. Nevoltris, D. de Bundel, S. Bossi, D. Moreno-Delgado, X. Rovira, T. C. Møller, D. El Moustaine, M. Mathieu, E. Blanc, H. McLean, E. Dupuis, G. Mathis, E. Trinquet, H. Daniel, E. Valjent, D. Baty, P. Chames, P. Rondard, J.-P. Pin, Allosteric nanobodies uncover a role of hippocampal mGlu2 receptor homodimers in contextual fear consolidation. Nat. Commun. 8, 1967 (2017).

17. Y. Ma, Y. Ding, X. Song, X. Ma, X. Li, N. Zhang, Y. Song, Y. Sun, Y. Shen, W. Zhong, L. A. Hu, Y. Ma, M.-Y. Zhang, Structure-guided discovery of a single-domain antibody agonist against human apelin receptor. Sci Adv. 6, eaax7379 (2020).

18. A. Wu, D. Salom, J. D. Hong, A. Tworak, K. Watanabe, E. Pardon, J. Steyaert, H. Kandori, K. Katayama, P. D. Kiser, K. Palczewski, Structural basis for the allosteric modulation of rhodopsin by nanobody binding to its extracellular domain. Nat. Commun. 14, 5209 (2023).

19. G. Corder, D. C. Castro, M. R. Bruchas, G. Scherrer, Endogenous and Exogenous Opioids in Pain. Annu. Rev. Neurosci. 41, 453–473 (2018).

20. B. L. Kieffer, C. J. Evans, Opioid receptors: from binding sites to visible molecules in vivo. Neuropharmacology. 56 **Suppl 1**, 205–212 (2009).

21. A. Manglik, H. Lin, D. K. Aryal, J. D. McCorvy, D. Dengler, G. Corder, A. Levit, R. C. Kling, V. Bernat, H. Hübner, X.-P. Huang, M. F. Sassano, P. M. Giguère, S. Löber, Da Duan, G. Scherrer, B. K. Kobilka, P. Gmeiner, B. L. Roth, B. K. Shoichet, Structure-based discovery of opioid analgesics with reduced side effects. Nature. 537, 185–190 (2016).

22. A. Faouzi, H. Wang, S. A. Zaidi, J. F. DiBerto, T. Che, Q. Qu, M. J. Robertson, M. K. Madasu, A. El Daibani, B. R. Varga, T. Zhang, C. Ruiz, S. Liu, J. Xu, K. Appourchaux, S. T. Slocum, S. O. Eans, M. D. Cameron, R. Al-Hasani, Y. X. Pan, B. L. Roth, J. P. McLaughlin, G. Skiniotis, V. Katritch, B. K. Kobilka, S. Majumdar, Structure-based design of bitopic ligands for the µ-opioid receptor. Nature. 613, 767–774 (2023).

23. H. Wang, F. Hetzer, W. Huang, Q. Qu, J. Meyerowitz, J. Kaindl, H. Hübner, G. Skiniotis, B. K. Kobilka, P. Gmeiner, Structure-based evolution of G protein-biased μ-opioid receptor agonists. Angew. Chem. Int. Ed Engl. 61, e202200269 (2022).

24. N. D. Volkow, F. S. Collins, The Role of Science in Addressing the Opioid Crisis. N. Engl. J. Med. 377, 391–394 (2017).

25. A. Manglik, A. C. Kruse, T. S. Kobilka, F. S. Thian, J. M. Mathiesen, R. K. Sunahara, L. Pardo, W. I. Weis, B. K. Kobilka, S. Granier, Crystal structure of the µ-opioid receptor bound to a morphinan antagonist. Nature. 485, 321–326 (2012).

26. Y. Zhuang, Y. Wang, B. He, X. He, X. E. Zhou, S. Guo, Q. Rao, J. Yang, J. Liu, Q. Zhou, X. Wang, M. Liu, W. Liu, X. Jiang, D. Yang, H. Jiang, J. Shen, K. Melcher, H. Chen, Y. Jiang, X. Cheng, M.-W. Wang, X. Xie, H. E. Xu, Molecular recognition of morphine and fentanyl by the human μ-opioid receptor. Cell. 185, 4361–4375.e19 (2022).

27. A. Koehl, H. Hu, S. Maeda, Y. Zhang, Q. Qu, J. M. Paggi, N. R. Latorraca, D. Hilger, R. Dawson, H. Matile, G. F. X. Schertler, S. Granier, W. I. Weis, R. O. Dror, A. Manglik, G. Skiniotis, B. K. Kobilka, Structure of the µ-opioid receptor-Gi protein complex. Nature. 558, 547–552 (2018).

28. Y. Wang, Y. Zhuang, J. F. DiBerto, X. E. Zhou, G. P. Schmitz, Q. Yuan, M. K. Jain, W. Liu, K. Melcher, Y. Jiang, B. L. Roth, H. E. Xu, Structures of the entire human opioid receptor family. Cell. 186, 413–427.e17 (2023).

29. A. J. S. Bloch, S. Mukherjee, J. Kowal, E. V. Filippova, M. Niederer, E. Pardon, J. Steyaert, A. Kossiakoff, K. P. Locher, Development of a universal nanobody-binding Fab module for fiducial-assisted cryo-EM studies of membrane proteins. Proc. Natl. Acad. Sci. U. S. A. 118 (2021), doi:10.1073/pnas.2115435118.

30. J. A. Ballesteros, H. Weinstein, "Integrated methods for the construction of three-dimensional models and computational probing of structure-function relations in G protein-coupled receptors" in Methods in Neurosciences, S. C. Sealfon, Ed. (Academic Press, 1995), vol. 25, pp. 366–428.

31. A. Manglik, Molecular Basis of Opioid Action: From Structures to New Leads. Biol. Psychiatry. 87, 6–14 (2020).

32. Z. Li, J. Liu, F. Dong, N. Chang, R. Huang, M. Xia, T. A. Patterson, H. Hong, Three-Dimensional Structural Insights Have Revealed the Distinct Binding Interactions of Agonists, Partial Agonists, and Antagonists with the µ Opioid Receptor. Int. J. Mol. Sci. 24 (2023), doi:10.3390/ijms24087042.

33. C. Chen, J. Yin, J. K. Riel, R. L. DesJarlais, L. F. Raveglia, J. Zhu, L. Y. Liu-Chen, Determination of the amino acid residue involved in [3H]beta-funaltrexamine covalent binding in the cloned rat mu-opioid receptor. J. Biol. Chem. 271, 21422–21429 (1996).

34. C. Marie-Pepin, S. Y. Yue, E. Roberts, C. Wahlestedt, P. Walker, Novel “restoration of function” mutagenesis strategy to identify amino acids of the δ-opioid receptor involved in ligand binding. J. Biol. Chem. 272, 9260–9267 (1997).

35. S. Granier, A. Manglik, A. C. Kruse, T. S. Kobilka, F. S. Thian, W. I. Weis, B. K. Kobilka, Structure of the δ-opioid receptor bound to naltrindole. Nature. 485, 400–404 (2012).

36. Y. Toyoda, A. Zhu, F. Kong, S. Shan, J. Zhao, N. Wang, X. Sun, L. Zhang, C. Yan, B. K. Kobilka, X. Liu, Structural basis of α1A-adrenergic receptor activation and recognition by an extracellular nanobody. Nat. Commun. 14, 1–13 (2023).

37. A. C. Hong, N. J. Byrne, B. Zamlynny, S. Tummala, L. Xiao, J. M. Shipman, A. T. Partridge, C. Minnick, M. J. Breslin, M. T. Rudd, S. J. Stachel, V. L. Rada, J. C. Kern, K. A. Armacost, S. A. Hollingsworth, J. A. O’Brien, D. L. Hall, T. P. McDonald, C. Strickland, A. Brooun, S. M. Soisson, K. Hollenstein, Structures of active-state orexin receptor 2 rationalize peptide and small-molecule agonist recognition and receptor activation. Nat. Commun. 12, 815 (2021).

38. Meredith A. Skiba, Sarah M. Sterling, Shaun Rawsona Morgan S.A. Gilman, Huixin Xu, Genevieve R. Nemeth, Joseph D. Hurley, Pengxiang Shen, Dean P. Staus, Jihee Kim, Conor McMahon, Maria K. Lehtinen, Laura M. Wingler, Andrew C. Kruse, Antibodies Expand the Scope of Angiotensin Receptor Pharmacology. BioRxiv (2023), doi:10.1101/2023.08.23.554128.

39. C. T. Dooley, N. N. Chung, P. W. Schiller, R. A. Houghten, Acetalins: opioid receptor antagonists determined through the use of synthetic peptide combinatorial libraries. Proc. Natl. Acad. Sci. U. S. A. 90, 10811–10815 (1993).

40. P. W. Schiller, G. Weltrowska, T. M.-D. Nguyen, C. Lemieux, N. N. Chung, Y. Lu, Conversion of δ-, κ- and μ-receptor selective opioid peptide agonists into δ-, κ- and μ- selective antagonists. Life Sci. 73, 691–698 (2003).

41. L. C. Purington, I. D. Pogozheva, J. R. Traynor, H. I. Mosberg, Pentapeptides Displaying μ Opioid Receptor Agonist and δ Opioid Receptor Partial Agonist/Antagonist Properties. J. Med. Chem. 52, 7724–7731 (2009).

42. P. V. Afonine, B. P. Klaholz, N. W. Moriarty, B. K. Poon, O. V. Sobolev, T. C. Terwilliger, P. D. Adams, A. Urzhumtsev, New tools for the analysis and validation of cryo-EM maps and atomic models. Acta Crystallogr D Struct Biol. 74, 814–840 (2018).

43. A. Punjani, J. L. Rubinstein, D. J. Fleet, M. A. Brubaker, cryoSPARC: algorithms for rapid unsupervised cryo-EM structure determination. Nat. Methods. 14, 290–296 (2017).

44. R. Sanchez-Garcia, J. Gomez-Blanco, A. Cuervo, J. M. Carazo, C. O. S. Sorzano, J. Vargas, DeepEMhancer: a deep learning solution for cryo-EM volume post-processing. Commun Biol. 4, 874 (2021).

45. P. Gouet, E. Courcelle, D. I. Stuart, F. Métoz, ESPript: analysis of multiple sequence alignments in PostScript. Bioinformatics. 15, 305–308 (1999).

46. J. Zivanov, T. Nakane, B. O. Forsberg, D. Kimanius, W. J. Hagen, E. Lindahl, S. H. Scheres, New tools for automated high-resolution cryo-EM structure determination in RELION-3. Elife. 7 (2018), doi:10.7554/eLife.42166.

47. S. Q. Zheng, E. Palovcak, J.-P. Armache, K. A. Verba, Y. Cheng, D. A. Agard, MotionCor2: anisotropic correction of beam-induced motion for improved cryo-electron microscopy. Nat. Methods. 14, 331–332 (2017).

48. A. Kucukelbir, F. J. Sigworth, H. D. Tagare, Quantifying the local resolution of cryo-EM density maps. Nat. Methods. 11, 63–65 (2014).

49. E. C. Meng, T. D. Goddard, E. F. Pettersen, G. S. Couch, Z. J. Pearson, J. H. Morris, T. E. Ferrin, UCSF ChimeraX: Tools for structure building and analysis. Protein Sci. 32, e4792 (2023).

50. A. Casañal, B. Lohkamp, P. Emsley, Current developments in Coot for macromolecular model building of Electron Cryo-microscopy and Crystallographic Data. Protein Sci. 29, 1069–1078 (2020).

51. D. Liebschner, P. V. Afonine, M. L. Baker, G. Bunkóczi, V. B. Chen, T. I. Croll, B. Hintze, L. W. Hung, S. Jain, A. J. McCoy, N. W. Moriarty, R. D. Oeffner, B. K. Poon, M. G. Prisant, R. J. Read, J. S. Richardson, D. C. Richardson, M. D. Sammito, O. V. Sobolev, D. H. Stockwell, T. C. Terwilliger, A. G. Urzhumtsev, L. L. Videau, C. J. Williams, P. D. Adams, Macromolecular structure determination using X-rays, neutrons and electrons: recent developments in Phenix. Acta Crystallogr D Struct Biol. 75, 861–877 (2019).

52. C. J. Williams, J. J. Headd, N. W. Moriarty, M. G. Prisant, L. L. Videau, L. N. Deis, V. Verma, D. A. Keedy, B. J. Hintze, V. B. Chen, S. Jain, S. M. Lewis, W. B. Arendall 3rd, J. Snoeyink, P. D. Adams, S. C. Lovell, J. S. Richardson, D. C. Richardson, MolProbity: More and better reference data for improved all-atom structure validation. Protein Sci. 27, 293– 315 (2018).

53. W. Kabsch, XDS. Acta Crystallogr. D Biol. Crystallogr. 66, 125–132 (2010).

54. J. Agirre, M. Atanasova, H. Bagdonas, C. B. Ballard, A. Baslé, J. Beilsten-Edmands, R. J. Borges, D. G. Brown, J. J. Burgos-Mármol, J. M. Berrisford, P. S. Bond, I. Caballero, L. Catapano, G. Chojnowski, A. G. Cook, K. D. Cowtan, T. I. Croll, J. É. Debreczeni, N. E. Devenish, E. J. Dodson, T. R. Drevon, P. Emsley, G. Evans, P. R. Evans, M. Fando, J. Foadi, L. Fuentes-Montero, E. F. Garman, M. Gerstel, R. J. Gildea, K. Hatti, M. L. Hekkelman, P. Heuser, S. W. Hoh, M. A. Hough, H. T. Jenkins, E. Jiménez, R. P. Joosten, R. M. Keegan, N. Keep, E. B. Krissinel, P. Kolenko, O. Kovalevskiy, V. S. Lamzin, D. M. Lawson, A. A. Lebedev, A. G. W. Leslie, B. Lohkamp, F. Long, M. Malý, A. J. McCoy, S. J. McNicholas, A. Medina, C. Millán, J. W. Murray, G. N. Murshudov, R. A. Nicholls, M. E. M. Noble, R. Oeffner, N. S. Pannu, J. M. Parkhurst, N. Pearce, J. Pereira, A. Perrakis, H. R. Powell, R. J. Read, D. J. Rigden, W. Rochira, M. Sammito, F. Sánchez Rodríguez, G. M. Sheldrick, K. L. Shelley, F. Simkovic, A. J. Simpkin, P. Skubak, E. Sobolev, R. A. Steiner, K. Stevenson, I. Tews, J. M. H. Thomas, A. Thorn, J. T. Valls, V. Uski, I. Usón, A. Vagin, S. Velankar, M. Vollmar, H. Walden, D. Waterman, K. S. Wilson, M. D. Winn, G. Winter, M. Wojdyr, K. Yamashita, The CCP4 suite: integrative software for macromolecular crystallography. Acta Crystallogr D Struct Biol. 79, 449–461 (2023).

55. A. J. McCoy, Solving structures of protein complexes by molecular replacement with Phaser. Acta Crystallogr. D Biol. Crystallogr. 63, 32–41 (2007).

56. T. Schwede, J. Kopp, N. Guex, M. C. Peitsch, SWISS-MODEL: An automated protein homology-modeling server. Nucleic Acids Res. 31, 3381–3385 (2003).

57. P. Emsley, K. Cowtan, Coot: model-building tools for molecular graphics. Acta Crystallogr. D Biol. Crystallogr. 60, 2126–2132 (2004).

